# A single-nucleus regulatory atlas links transposable elements to adult hippocampal neurogenesis

**DOI:** 10.64898/2026.08.31.748376

**Authors:** Umut Cakir, Federica Mantovani, Souad Youjil Abadi, Umer Javed Butt, Vikas Bansal, Riki Kawaguchi, Daniel Geschwind, Klaus-Armin Nave, Hannelore Ehrenreich, Manvendra Singh

## Abstract

Adult hippocampal neurogenesis can be enhanced or even triggered by extrinsic stimuli, but the lineage-specific regulatory changes that accompany a stimulated neurogenic response remain undefined. We used recombinant human erythropoietin (rhEPO), a defined influencing factor that enriches newly formed pyramidal neurons by ∼2 fold in this paradigm^1^, to profile ∼35,000 nuclei by single-nucleus ATAC sequencing (snATAC-seq) from rhEPO- and placebo-treated adult mouse hippocampi and to integrate them with matched single-nucleus RNA sequencing (snRNA-seq) of ∼120,000 nuclei. Thirty-five accessibility-defined clusters resolved 11 major lineages. Of 39,404 differentially accessible regions (DARs), more than half arose in pyramidal neurons, whereas dentate gyrus granule neurons and interneurons showed predominantly reduced accessibility; within the pyramidal lineage, remodelling was predominant in newly-formed rather than mature neurons. About 20% of hippocampal candidate cis-regulatory elements (cCREs) overlapped transposable elements (TEs) at baseline, rising to about 50% among regions that gained accessibility upon stimulation, and multiple families were significantly bound above the expected threshold by neurogenic transcription factors, whereby the many neighbouring genes have established roles for neuronal differentiation and synapse formation. These data define TE-derived sequences as candidate components of stimulus-responsive neurogenic regulatory networks and provide a cell-type-specific chromatin resource for the adult hippocampus at single-nucleus resolution.

**HIGHLIGHTS:**

- Single-nucleus atlas resolves chromatin accessibility across hippocampal lineages
- More than half of EPO-responsive accessible regions arise in pyramidal neurons
- Newly formed pyramidal neurons show the strongest chromatin remodelling
- TEs bound by neurogenic TFs serve as candidate cis-regulatory element for neurogenesis genes

**IN BRIEF:** Cakir et al. provide a lineage-resolved chromatin accessibility atlas of the adult mouse hippocampus under a controlled pro-neurogenic stimulus. Chromatin responses are unevenly distributed across lineages and concentrated in newly formed pyramidal neurons, and a substantial fraction of stimulus-responsive regions is derived from specific retrotransposons bound by neurogenic transcription factors.

## INTRODUCTION

Adult hippocampal neurogenesis is a stimulus-responsive process where the adult hippocampus retains progenitor populations whose differentiation can be modulated by physiological and pharmacological signals^2,3^. Most evidence of this responsiveness relied on cell counting, bulk expression profiling, or, more recently, single-cell transcriptomics^2–5^. What remains undefined is the regulatory layer that accompanies a stimulated neurogenic response, namely, which hippocampal lineages change their chromatin state, which candidate cis-regulatory elements (cCREs) open or compact, and which regulatory sequences supply the binding platforms for the transcription factors involved in adult neurogenesis. Because chromatin accessibility is measured at the level of individual loci, it offers a regulatory landscape of a stimulated hippocampus that transcript abundance alone cannot provide. Overlaying both the transcript abundances and chromatin landscape helps comprehending the landscape of regulatory changes, and the single nuclei resolution helps assigning these integrated layers in cell-type-, lineage-specific way.

Addressing this question requires an influencing factor that is defined, reproducible, dose-controlled, and already characterized at the transcriptional and physiological level, so that the chromatin data can be interpreted against a known cellular trajectory. Erythropoietin (EPO) satisfies these criteria. EPO is a hypoxia-inducible cytokine best known for its indispensable role in erythropoiesis, but it is also expressed by multiple cell types in the adult brain, with particularly high levels in the hippocampus, where it has been implicated as a pleiotropic neurotrophic factor that modulates synaptic plasticity, neuronal survival and circuit remodelling^1,6–8^. Recombinant human EPO (rhEPO) improves cognitive performance in several patient groups and confers neuroprotection in preclinical models of neurodegeneration and brain injury^9–11^, and several neuronal and glial lineages endogenously express EPO and its canonical receptor (EPOR)^12,13^, supporting an endogenous “neuro-EPO” system in the adult central nervous system^14^. Importantly for the present study, the cellular consequences of the rhEPO paradigm used here have already been established: our single-cell transcriptomic work showed that rhEPO promotes the development, differentiation and integration of pyramidal neurons - excitatory neurons of the hippocampus - while reducing the relative representation and inhibitory potential of GABAergic interneurons^1,15^, and electrophysiological recordings demonstrated enhanced excitatory and reduced inhibitory input onto newly integrated CA1 neurons^1^. rhEPO therefore functions here as an experimental tool with a known cellular outcome of unidirectional neurogenesis towards excitatory lineages.

However, the regulatory basis of that outcome is unresolved. Bulk genomics has shown that EPO treatment modulates plasticity-related gene expression and improves hippocampus-dependent behaviour, but lacks the resolution to connect these changes to defined cell types and cis-regulatory elements. EPO induces plasticity-associated genes and chromatin regulators, including *Bdnf*, *Gap43*, *Psd95*, *Egr1* and *Ep300*, and alters HDAC5 localization, pointing towards an epigenomic component^1,8,16^, yet whether and how any pro-neurogenic stimulus reshapes the chromatin landscape of specific hippocampal cell types *in vivo* has not been determined.

Single-nucleus assay for transposase-accessible chromatin sequencing (snATAC-seq) provides genome-wide accessibility profiles at single-cell resolution and thereby identifies cCREs, the promoters and distal enhancers that underpin cell-type-specific transcriptional programs^17^. While the reference atlases of the adult mouse brain have catalogued such elements in the unperturbed state^18–20^, stimulated adult hippocampus has not been profiled yet. Applying snATAC-seq to rhEPO- and placebo-treated hippocampi allows regulatory events to be captured in rare subpopulations, including the newly formed neurons that are the outcome of ongoing neurogenesis^1,6^.

An unresolved question concerns the origin of the regulatory elements that fuel adult neurogenesis in response to stimuli. Roughly half of the mouse and human genome consists of transposable elements (TEs), including LINEs, SINEs, endogenous retroviruses (ERVs) and DNA transposons^21,22^. TEs have been repeatedly co-opted into gene regulatory networks in development, immunity and also in the brain^23,24^; a substantial subset of brain-specific cCREs is TE-derived, and in the human cortex approximately 80% of recently gained, human-specific regulatory elements overlap with TEs^25^. ERVs, SINEs and DNA transposons have been shown to serve as cCREs in neural progenitors, supplying novel binding sites for neurogenic transcription factors^26,27^. Whether TE-derived sequences form part of the regulatory repertoire that responds to a neurogenic stimulus in the adult hippocampus remains to be tested.

Here we combine snATAC-seq with matched snRNA-seq from the same animals and with published transcription-factor ChIP-seq datasets to describe the regulatory landscape of the adult mouse hippocampus under sustained rhEPO treatment. We first construct an integrated single-nucleus atlas of chromatin accessibility and gene expression across major hippocampal lineages under placebo and rhEPO conditions. We then define treatment-responsive differentially accessible regions (DARs) globally and lineage-by-lineage, relate them to gene expression, transcription-factor motif usage and inferred gene regulatory networks (GRNs), and resolve them further within newly formed versus mature pyramidal neurons. Finally, we quantify the contribution of TE-derived sequences to stimulus-responsive cCREs and ask whether these sequences are preferentially occupied by neurogenic transcription factors in independent ChIP-seq data in the vicinity of neurogenesis- or synaptic-related genes.

The resulting atlas shows the known and new accessible regions in the adult hippocampus. The chromatin response to a defined neurogenic stimulus is not evenly distributed across hippocampal lineages but is concentrated in newly formed excitatory neurons. The chromatin accessibility changes are concordant with, and largely proximal to, transcriptional changes in the same cells, and selected TEs comprise around half of the stimulus-responsive regions. These TEs are embedded with and are bound by neurogenic transcription factors far above the expected chance. We frame these TE-derived regions as cCREs elements for a group of close-by genes with established roles in neurogenesis and synapse formation. The accessibility, transcription-factor occupancy and association with the expression of neighbouring differentiation and synapse genes are associative, and direct influence would be required to establish regulatory function. Beyond these observations, the dataset is released as a resource for examining stimulus-responsive regulatory landscapes and cell-type-specific cCREs in the adult brain.

## RESULTS

### An integrated single-nucleus multiomics atlas of the adult hippocampus resolves 35 clusters

To resolve chromatin accessibility in the adult hippocampus at the level of individual cell types, and to do so under a controlled pro-neurogenic stimulus, we performed snATAC-seq on hippocampi of adult C57BL/6N mice treated with an established dosing paradigm (11 injections every other day from P28 to P49)^1^. Eight libraries were generated (4 rhEPO, 4 placebo), each prepared from left hippocampi pooled from two mice of the same treatment group, with the matched snRNA-seq dataset generated from the pooled right hippocampi of the same animals (GSE220522)^1^. Libraries were prepared on the 10x Genomics Chromium platform and processed with Cell Ranger ATAC^28^. After quality control on fragment number, TSS enrichment, nucleosome signal and blacklist fraction, ∼35,000 high-quality nuclei were retained; per-sample nucleus counts are given in Table S1 (Extended Data Fig. S1A-B). The libraries showed strong TSS enrichment and clear nucleosomal banding, with <5% of fragments mapping to ENCODE blacklist regions (Extended Data Fig. S2A-B).

We constructed a unified peak set across samples in Signac^29^, applied TF-IDF normalization and latent semantic indexing, and integrated samples with Harmony to correct batch effects^30^ (Fig. 1A). Graph-based clustering resolved 35 accessibility-defined clusters (Fig. 1B and Table S2). Cell identities were assigned by integrating the snATAC-seq data with the matched snRNA-seq profiles; anchoring ATAC-derived gene activity scores to RNA expression by canonical correlation analysis produced a joint embedding in which the two modalities overlapped (Fig. 1C), and label transfer yielded high-confidence annotations (Fig. 1D-E). Eleven major hippocampal lineages were distinguished: dentate gyrus granule neurons, excitatory (pyramidal) neurons, interneurons, microglia, astrocytes, oligodendrocytes, endothelial cells, pericytes, intermediate progenitors, ependymal cells and neuroimmune cells (Fig. 1B-C and Table S3). Accessibility at canonical marker loci was consistent with these assignments^19^, with astrocyte clusters accessible at *Slc1a2*, oligodendrocyte-lineage clusters at *Plp1*, *Mbp*, *Mog* or *Pdgfra*, dentate gyrus neurons at *Prox1*, and excitatory neurons at *Slc17a7* (Fig. 1G and Tables S2-S4), and chromatin-defined identities corresponded to the transcriptomic clusters of the companion snRNA-seq dataset (Fig. 1C-E and Extended Data Fig. S3; Fisher’s exact test, p < 0.05). Together, this integrated multi-omic atlas delineates the cell-type-specific regulatory landscape of the adult hippocampus, linking cCREs to gene expression programs and further highlights distinct cis-regulatory signatures for each cell type (Fig. 1F).

**Figure 1.**
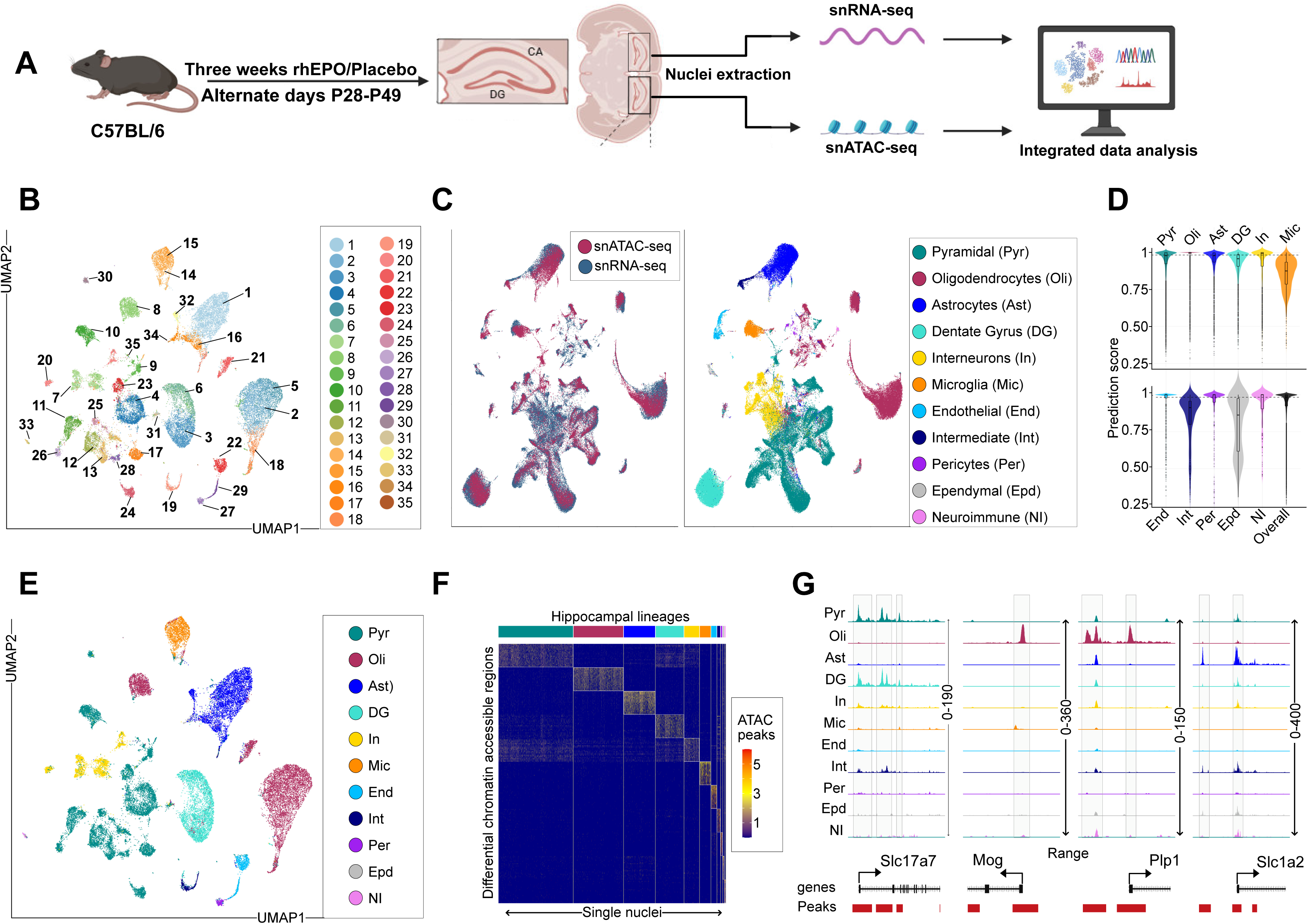
An integrated single-nucleus chromatin accessibility and transcriptome atlas of the adult hippocampus under a defined pro-neurogenic stimulus. **(A)** C57BL/6N mice received recombinant human erythropoietin (rhEPO) or placebo every other day from postnatal day (P) 28 to P49. Hippocampal regions were dissected and nuclei isolated for single-nucleus RNA-seq (snRNA-seq) and single-nucleus ATAC-seq (snATAC-seq). The resulting datasets were jointly analyzed to integrate gene expression and chromatin accessibility. Data processing included Cell Ranger peak calling, Signac-based quality control and preprocessing, Harmony batch correction, co-embedding of snRNA-seq and snATAC-seq, cell-type annotation, differential accessibility testing, motif enrichment, gene regulatory network reconstruction, and identification of gene- and transposable element (TE)-associated candidate cis-regulatory elements (cCREs). **(B)** UMAP projection of the snATAC-seq dataset coloured by 35 unbiased Seurat clusters. **(C)** Joint embedding of snATAC-seq and snRNA-seq using gene activity and expression. Left: UMAP projection of integrated nuclei, with snATAC-seq (red) and snRNA-seq (blue) showing strong overlap following CCA-based integration of ATAC-derived gene activity scores and RNA expression. Right: the same embedding coloured by major cell types assigned via label transfer and canonical markers, including pyramidal neurons (Pyr), dentate gyrus neurons (DG), interneurons (In), oligodendrocytes (Oli), astrocytes (Ast), microglia (Mic), endothelial cells (End), intermediate progenitors (Int), pericytes (Per), ependymal cells (Epd) and neuroimmune cells (NI). **(D)** Violin plots of prediction scores for cell-type assignments obtained by label transfer from snRNA-seq to snATAC-seq. Each violin represents a major hippocampal cell type; the final column in the lower panel shows the overall distribution across all types. The dashed line marks the median overall prediction score. **(E)** UMAP projection of the snATAC-seq dataset, coloured by cell-type identities transferred from the matched snRNA-seq reference shown in (C). **(F)** Heatmap of differentially accessible chromatin regions (DARs, rows) across hippocampal cell types (columns). Scaled accessibility values (red/yellow = high, blue = low) indicate distinct cis-regulatory signatures for each lineage. For visualization, the top 20 peaks per cell type are shown. **(G)** Genome browser tracks showing chromatin accessibility at the Slc17a7, Mog, Plp1 and Slc1a2 loci across major hippocampal cell types. Highlighted regions mark cell-type-restricted accessible regions, illustrating lineage-specific regulatory landscapes.

Peak distribution differed between lineages, indicating distinct cis-regulatory repertoires per cell type (Fig. 1F). These integrated multi-omic analyses provide a foundation for constructing gene regulatory networks underlying hippocampal cell identity. For instance, we show here one locus that illustrates the resolution the atlas provides: the *Slc1a2* (*EAAT2*) promoter is broadly accessible across hippocampal lineages, whereas its distal elements are open specifically in astrocytes, matching astrocyte-specific expression (Fig. 1G). This is consistent with the general observation that promoters are frequently accessible across cell types while enhancer deployment is cell-type-restricted^31,32^. The atlas therefore, provides a cell-type-resolved map of cCREs in the adult hippocampus and the framework for all subsequent comparisons between treatment conditions.

### Hippocampal cCREs encode lineage-specific TF motifs and a substantial TE contribution

We first asked which regulatory codes and which sequence classes constitute this landscape, independently of treatment. To obtain an overview of regulatory diversity, we identified cell-type-specific DARs across the 11 lineages and subjected to motif enrichment analysis. We detected 135 TF motifs significantly enriched in cluster- and lineage-associated peaks (log2 enrichment ratio > 2 and adjusted p < 1 x 10^-10^; Fig. 2A, Extended Data Fig. S4, Tables S2-S3). As expected, well-known neurogenic and lineage-specific factors were prominent, including SOX factors^33^, EGR family members^34^, the NEUROD1/2 and NEUROG2 factors essential for neuronal differentiation^35^, and NRF1, which activates neuronal and synaptic genes^36^. We also identified motifs for neurodevelopmental regulators less emphasized in hippocampal biology, but linked to neurodevelopment, including GLIS1/2, ZBTB14, RFX2/5, the NR4A2::RXRA heterodimer and ZNF423^37^. Enrichment was lineage-patterned: OLIG-like and SOX-family motifs predominated in glial clusters, consistent with oligodendrocyte and astrocyte programs^38^; EGR/ERG motifs dominated progenitor and early neuronal clusters but not interneurons, reflecting excitability-dependent genes and cell-cycle control in neural precursors^34^; and ETS/ELF motifs were enriched in endothelial and microglial clusters, aligning with their immune regulatory programs^20^ (Fig. 2A, Extended Data Fig. S4, Tables S2-S3). This integrated atlas, therefore, captures both the cellular composition and the underlying regulatory codes of the adult hippocampus regardless of the treatment.

**Figure 2.**
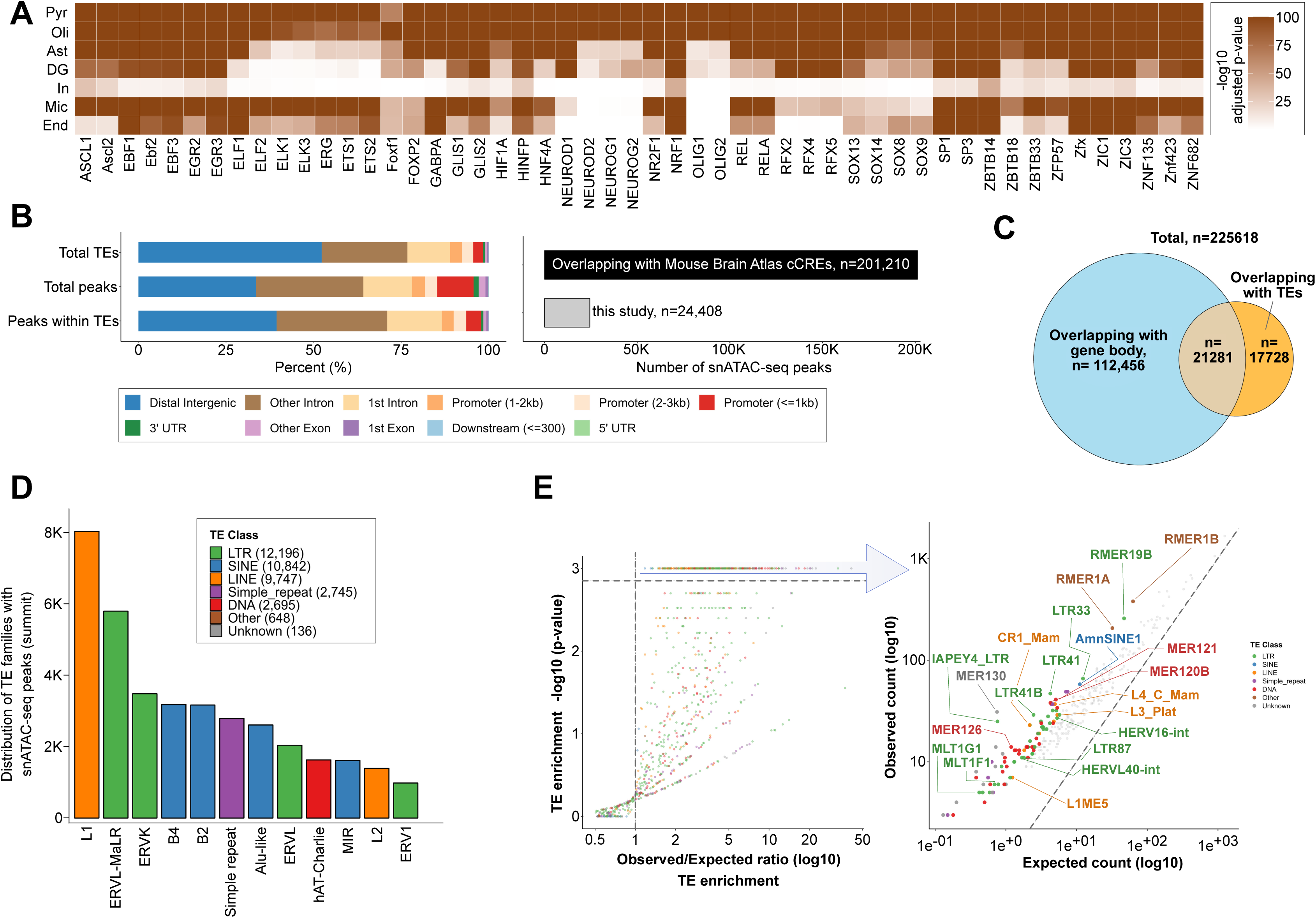
Cell-type motif usage and the sequence composition of hippocampal candidate cis-regulatory elements. **(A)** Heatmap of significantly enriched transcription factor (TF) motifs in cell-type-specific DARs. Rows correspond to 7 major hippocampal cell types and columns to representative motifs (colour scale = -log10 adjusted p-value). The remaining 4 cell types are not shown owing to limited motif detection resulting from their comparatively lower cell numbers. **(B)** Genomic distribution of (i) all annotated TEs in the mouse genome, (ii) all snATAC-seq peaks identified in this study (225,618 summits) and (iii) the subset of peaks overlapping TE sequences (39,009 summits). Bars indicate the percentage of features located in distal intergenic, intronic, exonic, promoter (<1 kb, 1-2 kb or 2-3 kb), downstream or UTR regions. Right: overlap of snATAC-seq peaks with CATlas, showing 201,210 overlapping peaks (black) and 24,408 non-overlapping peaks (grey). **(C)** Venn diagram summarising the intersection between peaks that map to gene bodies (112,456), TEs (17,728) or both (21,281). **(D)** Bar chart of the TE families at peak summits, coloured by TE class (LTR, SINE, LINE, simple repeat, DNA, other and unknown). **(E)** Left: log2(observed/expected) overlap for each TE family versus -log10 p-value. Right: observed versus expected counts for the top significantly enriched families highlighted at left.

Upon annotating snATAC-seq peaks by genomic context, we noticed the most accessible regions mapped outside annotated promoters. In the genomic context, ∼10.4% of accessible regions overlapped annotated promoters (+/-1 kb from TSS), whereas 44.5% fell within introns, 33.5% in intergenic regions and 4.3% in exons or UTRs (Fig. 2B, Extended Data Fig. S5, Table S4), a distribution consistent with most cCREs residing in intronic and intergenic space. Comparison with a published single-cell atlas of adult mouse brain chromatin accessibility^19,20^ showed that ∼10.8% of our hippocampal peaks were absent from prior annotations, defining a set of previously unannotated, potentially hippocampus-enriched cCREs (Fig. 2B, Extended Data Fig. S5A, Table S4). Because TEs constitute roughly half of the mammalian genome^22^ and contribute substantially to regulatory landscapes^39^, we intersected peaks with RepeatMasker annotations to measure the extent to which TEs contribute to this landscape. Approximately 20% of hippocampal cCREs overlapped annotated TEs, spanning LTR, LINE, SINE and other classes (Fig. 2C-E, Extended Data Fig. S5B-C, Table S5), close to the ∼25% reported for mammalian cCREs^39^. Notably, LTR elements exhibited disproportionately high accessibility, supporting the view that LTRs often function as self-contained regulatory modules for neighbour gene expression^37,39^. These TE-derived cCREs may serve as lineage-specific regulatory elements influencing hippocampal gene programs^40,41^. Together, these findings reveal that TEs are extensively integrated into the adult hippocampal regulatory network, providing a pool of fresh cell-type-enriched cCREs that likely contribute to hippocampal function.

### The global chromatin response to stimulation is enriched for neurogenic genes and TE-derived regions

We first asked how the stimulus affects hippocampal chromatin when cell identity is disregarded, comparing aggregated peaks between rhEPO- and placebo-treated samples. Logistic regression-based differential accessibility analysis identified 945 DARs (FDR < 0.01), of which 799 gained and 146 lost accessibility under rhEPO (Fig. 3A). Of these, 639 were located within +/-1 kb of an annotated TSS, and around one-third of the fraction (n=231) overlapped TE sequences (Fig. 3B). Gene ontology analysis of the associated genes returned enrichment for neuroplasticity, synapse organization and cognitive function (Fig. 3C, Table S6). Individual loci included the *Ascl1* promoter, whose product promotes neural progenitor differentiation^42^, and *Egr2*, an immediate-early gene of the EGR family implicated in neuronal plasticity^43^ (Fig. 3A).

**Figure 3.**
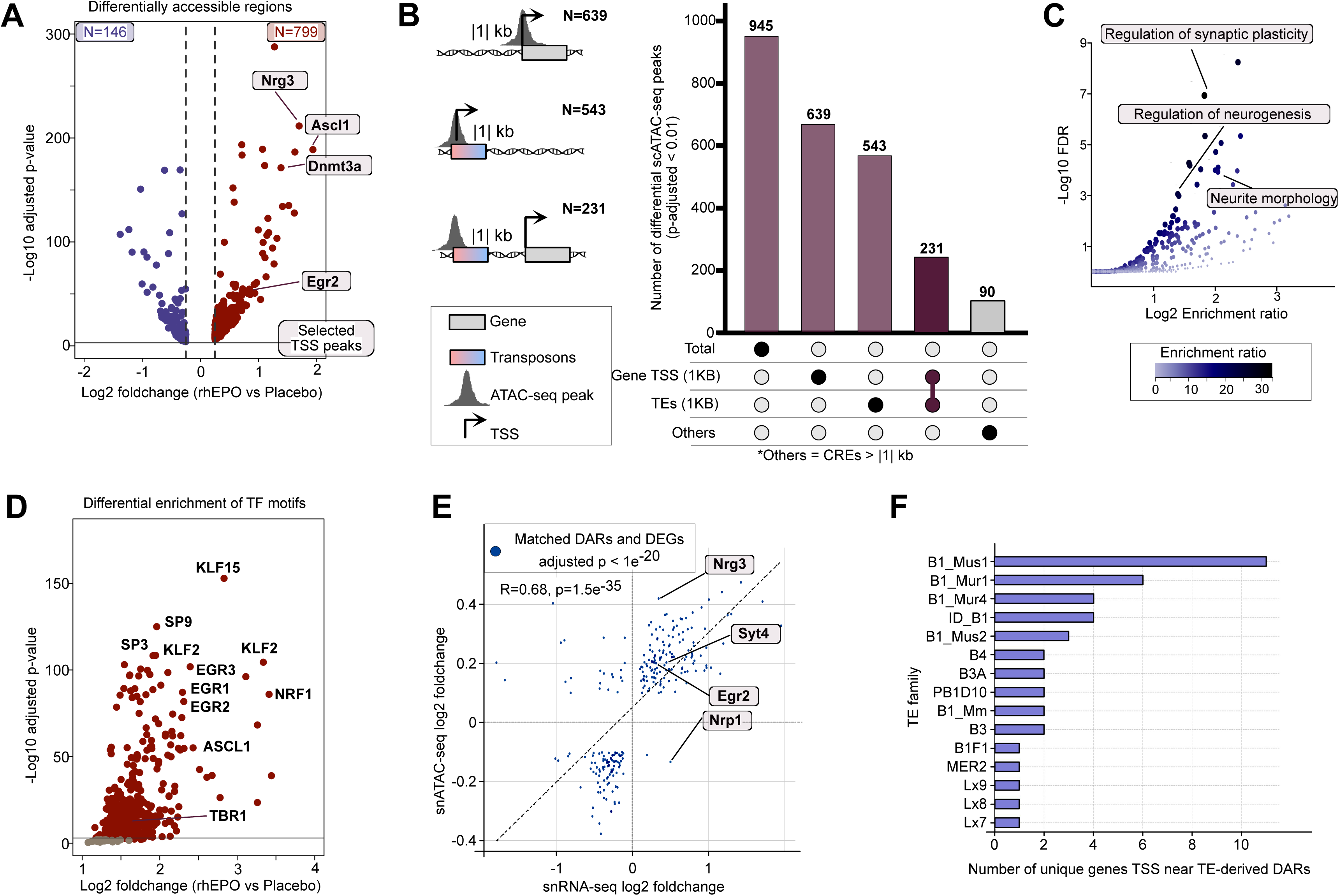
The global chromatin response to stimulation is numerically restricted and enriched for TE-derived regions. **(A)** Volcano plot of DARs between rhEPO- and placebo-treated hippocampi, identifying 945 DARs (799 gained, 146 lost; FDR < 0.01). Neurogenic regulators (Ascl1, Dnmt3a, Nrg3, Egr2) are highlighted. **(B)** DARs grouped into three categories: promoter-proximal (within +/-1 kb of a TSS), TE-associated (DARs overlapping TEs within +/-1 kb of a TSS) and distal cCREs (>1 kb from a TSS). The bar plot and accompanying UpSet diagram summarise the counts per category, with schematic illustrations at left indicating classification criteria. Filter applied here is based on -log10p-values > 2. **(C)** Gene ontology enrichment of DAR-associated genes, with significant terms related to synaptic plasticity, neurogenesis and neurite morphology. **(D)** Motif enrichment analysis of DARs, showing over-representation of TFs including KLFs, EGR1-3, ASCL1, NRF1, SP-family factors and TBR1. **(E)** Scatter plot of gene activity changes inferred from snATAC-seq versus differentially expressed genes (DEGs) from snRNA-seq. Log2 fold changes (rhEPO versus placebo) are concordant (R = 0.68, p = 1.5 x 10-35), with representative neurogenic genes (Nrg3, Syt4, Egr2, Nrp1) highlighted. **(F)** Number of unique genes with a TSS located within +/-1 kb of TE-derived DARs. Rodent-specific B1 SINE subfamilies (B1_Mus1, B1_Mur1, B1_Mur4, ID_B1) are the most frequent contributors, linking TE-derived accessibility to candidate gene regulation.

Motif enrichment within these DARs belonged to families associated with neurogenesis and synaptic plasticity (Fig. 3D). Beyond the enhancer-associated SP1/2 motifs, rhEPO-accessible regions were enriched for Kruppel-like factor (KLF) motifs, consistent with KLF involvement in neuronal maturation and adult neurogenesis^44^, and for early growth response factors (EGR1-3), immediate-early TFs critical for learning and memory^34,45^. Motifs for AP-2 (TFAP2) factors, CREB/ATF members including Atf1, and GLIS factors were also over-represented (Table S5); these families have described roles in neuronal maturation, learning and memory, and activity-dependent gene expression^46^. Thus, at a global level, EPO promotes chromatin opening at promoter-proximal and distal sites that harbour motifs for neurogenic and plasticity-associated TFs, and often overlap with TE sequences.

We next overlaid and compared our snATAC-seq results with snRNA-seq and tested the levels of concordance between them. Comparison with differentially expressed genes (DEGs) from the matched snRNA-seq dataset showed that accessibility and expression changes were concordant, particularly for DARs proximal to DEGs (R = 0.68, p = 1.5 x 10^-35^; Fig. 3E). *Egr2*, *Syt4*, *Nrg3*, *Nrp1* and further DAR-associated genes were expressed at higher levels in rhEPO hippocampi (Fig. 3E, Table S7)^1,6^. Several synaptic loci showed increased accessibility only in rhEPO nuclei, including *Syt4*, an activity-inducible vesicle protein that modulates BDNF release and learning-related plasticity, and *Nrg3*, a trophic factor that promotes excitatory synapse formation on interneurons; differential peaks were also present at the *Nrp1* locus, encoding a receptor for semaphorin axon guidance cues^47^. Because both modalities were sampled at a single time point after treatment, these data establish concordance between the chromatin and transcriptional layers but do not order them temporally, thus suggesting a coordinated epigenome-transcriptome response underlying EPO-induced alterations in the neural network.

Strikingly, approximately 50% of regions gaining accessibility overlapped TEs, compared with ∼20% across the hippocampal cCRE repertoire. These were predominantly rodent-specific SINEs of the B1/Alu and B2 families, including *ID_B1*, *B3* and *B1_Mus1/2* subtypes (Fig. 3B, 3F), consistent with reports that SINE-derived DNA and transcripts can serve normal neuronal functions, and that SINE retrotransposons across the mammalian clade can recruit transcriptional regulators or RNA-binding proteins to nearby genes in *cis*^48,49^. The proximity of enriched B1/B2 elements to DEG-associated loci (Fig. 3F) identifies these sequences as candidate contributors to the regulatory response; the analysis is an overlap enrichment and does not establish that any individual element is functional. Overall, our analysis shows that the genome-wide chromatin changes upon neurogenic stimulus preferentially target a set of of TE-derived loci belonging to retrotransposon sequences that may be recruited to mediate the regulation of gene programs.

### rhEPO remodelled chromatin architecture is predominantly enriched in pyramidal neurons

We next asked whether the response is distributed evenly across hippocampal cell types or restricted to particular lineages. To determine, we tested differential accessibility within each of the 11 major lineages as defined earlier^1^ from the same animals. Using a logistic regression test at adjusted p < 0.01, we identified 39,404 DARs across all lineages. Of these, 7,608 were shared by all lineages and 21,555 were specific to a single lineage, with the remainder shared by more than one lineage but not by all (Fig. 4A, Table S8). Pyramidal neurons alone accounted for more than half of all DARs (Fig. 4B, Table S8).

**Figure 4.**
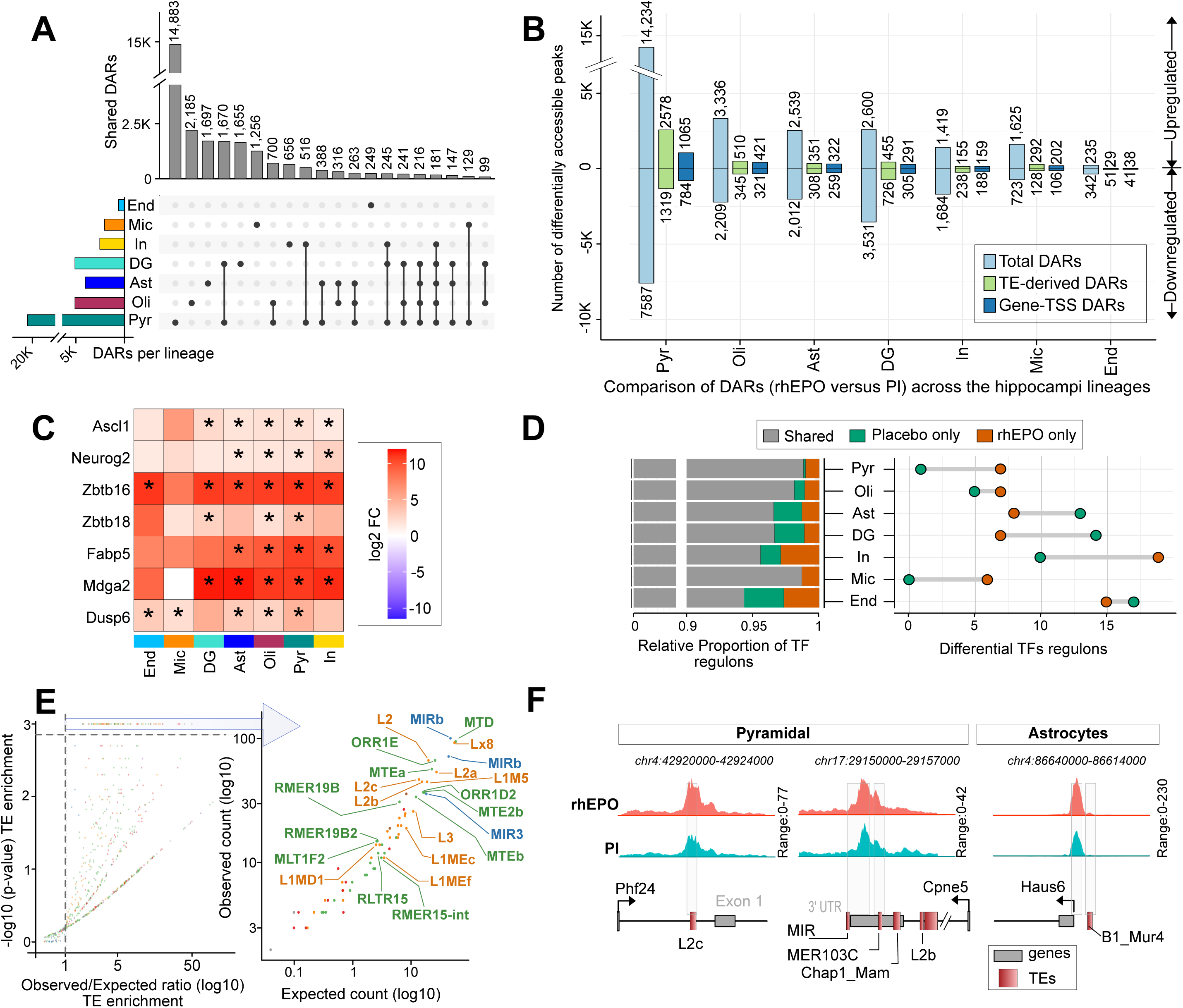
Chromatin remodeling, transposon usage and TF programs are lineage-selective across hippocampal cell types. **(A)** Global view of DARs (adjusted p < 0.01) between rhEPO- and placebo-treated nuclei. Horizontal bars indicate the number of DARs identified in each lineage (colour legend, left); the UpSet matrix at right shows how many DARs are unique to one lineage, shared by several, or shared by all. **(B)** Stacked bars show, for each hippocampal lineage, the total number of DARs (light blue), the subset overlapping TEs (green) and the subset located within +/-1 kb of gene TSSs (dark blue). Bars extending upward represent DARs with increased accessibility in rhEPO-treated samples; downward bars indicate reduced accessibility relative to placebo. **(C)** Heatmap of rhEPO-associated promoter accessibility changes at selected neurogenic genes across hippocampal lineages. Colours indicate log2 fold change (rhEPO versus placebo), with red showing increased and blue decreased accessibility. Asterisks denote significant changes (FDR < 0.01). **(D)** Stacked bars show the proportion of regulons inferred by Pando that are shared, rhEPO-specific or placebo-specific within each lineage. The dot plot at right summarises the absolute number of differential regulons, with orange indicating rhEPO-specific and green placebo-specific regulons. **(E)** TE enrichment analysis for rhEPO-responsive DARs. Left: log2(observed/expected) overlap versus -log10 P for every annotated TE family (points coloured by TE class; dashed boxes mark the top significantly enriched families). Right: observed versus expected counts for the top significant families. **(F)** Genome browser snapshots of rhEPO-associated chromatin accessibility at TE-derived candidate regulatory elements. In pyramidal neurons, increased accessibility was observed at an L2c element near Phf24 and at MIR and MER103C elements near Cpne5. In astrocytes, a B1_Mur4 element near Haus6 showed enhanced accessibility. Tracks display snATAC-seq signal for rhEPO (red) and placebo (blue), with genes (grey) and TEs (red) shown below.

The number of promoter-proximal and TE-overlapping DARs generally reflected the overall number of DARs in each cell type. Pyramidal neurons contained the largest numbers of both (Fig. 4B, Extended Data Fig. S6A, Table S8). Dentate gyrus granule neurons contained the next highest DAR count (n = 6,131) but showed the opposite direction. Most TE-associated regions lost accessibility (455 gained versus 726 lost), as did promoter regions (291 gained versus 305 lost). Oligodendrocytes (742 DARs), astrocytes (581 DARs) and microglia (308 DARs) showed moderate responses with a slight bias toward increased accessibility. Interneurons contained 347 DARs, of which 159 gained and 188 lost accessibility. Endothelial cells, pericytes, neuroimmune cells, intermediate progenitors and ependymal cells showed the fewest changes (Fig. 4B, Extended Data Fig. S6A, Table S8).

The chromatin response to this stimulus is therefore lineage-selective in both magnitude and direction: gains are concentrated in pyramidal neurons and, to a lesser degree, glial populations, whereas dentate gyrus granule neurons and interneurons show a net loss of accessibility. This directional split parallels the shift in cellular composition previously reported for the same paradigm^1,15^, although the present data cannot distinguish whether the reduced accessibility in inhibitory and granule lineages reflects regulatory change within persisting cells, or changes in the representation of cells recovered from each lineage.

We next asked if the key neurogenic gene promoter accessibilities are significantly upregulated in rhEPO samples. Consistent with the directional pattern in excitatory and glial lineages, promoters of neurogenic regulators showed accessibility gains. Expectedly, pyramidal neurons showed the activation of key neurogenic TFs and developmental regulators: *Ascl1*, *Neurog2* and *Pax6*, key factors driving neuronal differentiation from precursors, and *Nrg3*, a neuregulin promoting excitatory synaptogenesis^50^, all exhibited robust accessibility gains in pyramidal neurons, followed by dentate gyrus, astrocytes, and oligodendrocytes (Fig. 4C, Extended Data Fig. S6B, Table S9). Less-characterised regulators also emerged among the top stimulus-responsive promoters, including the zinc-finger factors *Zbtb16* (Plzf) and *Zbtb18* (Rp58), both implicated in cortical neurogenesis and neuronal subtype development^50^, together with *Fabp5*, *Mdga2* and *Dusp6* (Fig. 4C, Extended Data Fig. S6B, Table S9). The enrichment of these diverse promoters suggests that there is a broad activation of pro-neurogenic chromatin landscape upon rhEPO treatment, encompassing classical neurogenesis drivers and new regulatory elements linked to neuronal maturation.

### Neurogenic TFs and specific TE families are enriched within stimulus-responsive regulatory networks

Following these leads, we asked what differentially active Gene Regulatory Networks (GRNs) in hippocampal pyramidal and glial lineages respond to the stimulus. To relate accessibility changes to network structure, we applied the Pando algorithm to the matched snRNA-seq and snATAC-seq datasets, identifying transcription factors, their target regulatory regions and downstream effector genes^51^. The inferred networks are computational predictions from co-accessibility and co-expression, demonstrating the regulatory relationships between a group of accessible regions and their neighbour gene expression as distinct “regulons”. Most regulons were shared between rhEPO and placebo conditions; approximately a dozen were enriched under rhEPO (Fig. 4D, Extended Data Fig. S7, Table S10), and these were driven predominantly by transcription factors involved in neuronal differentiation and maturation.

A broad spectrum of TE families was significantly enriched within the DARs (Fig. 4E, Extended Data Fig. S8, Table S11), including LTR retrotransposons, namely ORR1E and MTD families, the L2/L2a LINEs, the rodent SINE families B3 and B4/B4A, and the MIR/MIRb elements^49,52,53^. Such elements have already been described as co-opted cCREs in neural cells^54^. Individual examples of increased accessibility under rhEPO included L2c, MER103C/MIR and B1_Mur4 elements located near *Phf24*, *Cpne5* and *Haus6*, respectively (Fig. 4F). These elements are candidate cis-regulators of the neighbouring genes. These patterns indicate that hippocampal stimulation unlocks repeat-rich genomic regions in a lineage-specific manner, suggesting these TEs act as cCREs that influence the transcription of nearby genes.

### Newly formed pyramidal neurons show the largest chromatin response to rhEPO

Given that pyramidal neurons carry more than half of all DARs, we asked whether the response is uniform across this lineage or restricted to a maturation stage. Excitatory neurons, particularly CA1 pyramidal cells, were previously reported to be over-represented and GABAergic interneurons under-represented^1,15^. In the present chromatin dataset, newly formed pyramidal subpopulations were expanded up to 5-fold (∼500% of control levels) in rhEPO samples (Extended Data Fig. S9A-B). This exceeds the ∼2-fold (∼200%) enrichment of immature pyramidal neurons reported from single-cell transcriptomic analysis of the same paradigm, and the ∼20% increase in mature CA1 pyramidal neuron numbers reported after prolonged treatment^1,6^; a single EPO dose has been reported to acutely elevate immature glutamatergic precursors within hours^6^. Because the present measurement is a proportional abundance derived from nuclei recovered for chromatin profiling, and because it disagrees quantitatively with the transcriptomic estimate obtained under the same regimen, we treat the ∼200% value as the reference estimate and the chromatin-derived value as an upper bound (*see Discussion*).

To resolve the lineage further, we clustered pyramidal neurons at high resolution in the snATAC-seq data. Initial clustering of the snRNA-seq data had identified 20 pyramidal clusters^1^, whereas snATAC-seq resolved 29 (Extended Data Fig. S10A). Cross-modal integration annotated these into 17 distinct cell types with substantial concordance between modalities (Fig. 5A, Extended Data Fig. S10B-D). The two predominant subgroups, newly formed neurons (early-stage markers) and mature neurons (late-stage markers), segregated into discrete UMAP clusters and were congruent between modalities. Comparing these two populations directly, we identified 7,420 DARs distinguishing newly formed from mature pyramidal neurons, of which 1,705 mapped within gene promoters and 935 overlapped TE sequences (Fig. 5B, Table S12). Approximately 66% of promoter-associated DARs were more accessible in newly formed neurons, and gene ontology analysis of the associated genes returned enrichment for neurogenesis and synapse formation (Fig. 5C). Of note, among DARs that were both promoter-associated and TE-overlapping, ∼83% showed higher accessibility in rhEPO samples.

**Figure 5.**
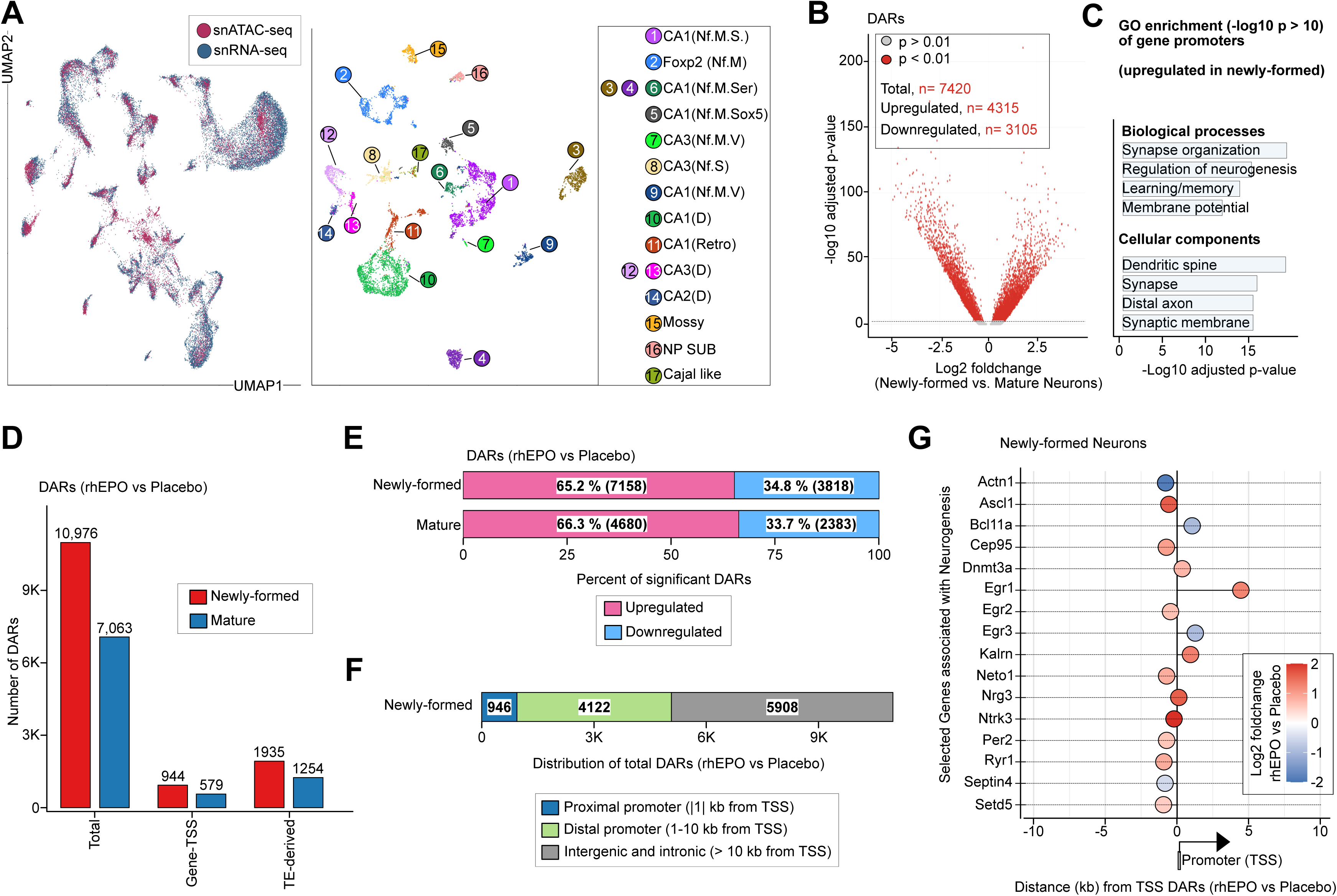
Within the pyramidal lineage, chromatin remodeling is concentrated in newly formed neurons. **(A)** Left: joint UMAP embedding of pyramidal-lineage nuclei profiled by snATAC-seq (red) and snRNA-seq (blue). Right: UMAP of the same nuclei coloured by clusters annotated through label transfer and marker gene activity, resolving newly formed neurons, mature pyramidal neurons and other subtypes (for example mossy, NP SUB and Cajal-like cells). **(B)** Volcano plot of DARs between newly formed and mature pyramidal neurons (FDR < 0.01). A total of 7,420 DARs were identified, including 4,315 with increased and 3,105 with reduced accessibility in newly formed neurons. **(C)** Gene ontology enrichment analysis of promoter DARs with increased accessibility in newly formed neurons. Significant terms (-log10 p > 10) include synapse organization, neurogenesis, learning, memory and membrane potential, and neuronal compartments (dendrite, synapse, distal axon and synaptic membrane). **(D)** Numbers of DARs in newly formed and mature neurons (rhEPO versus placebo), shown for all DARs (left), promoter-proximal gene-TSS DARs (middle) and TE-derived DARs (right). Newly formed neurons exhibit higher counts across all categories. **(E)** Proportional distribution of the DARs in (D). In both newly formed and mature neurons the majority of DARs are increased in accessibility (pink) and a smaller fraction reduced (blue). **(F)** Genomic distribution of rhEPO-responsive DARs in newly formed neurons relative to the nearest TSS. DARs are classified as promoter-proximal (<=1 kb, blue), distal promoter (1-10 kb, green) or intergenic/intronic (>10 kb, grey). **(G)** Distance of significant DAR summits to the nearest TSS for selected neurogenic and synaptic genes in newly formed neurons. Each dot represents the nearest DAR to a gene TSS, coloured by log2 fold change (red = increased, blue = decreased accessibility).

Comparing rhEPO with placebo within each population, mature pyramidal neurons showed ∼7,000 DARs and newly formed neurons ∼11,000, with ∼65% gained and ∼35% lost in both cases; exact counts are given in Table S13 (Fig. 5D-E, Extended Data Fig. S11A). Within newly formed neurons, 946 DARs were promoter-proximal (+/-1 kb from TSS) and a further ∼4,000 were distal (within +/-10 kb of a TSS), marking candidate enhancers and distal cCREs (Fig. 5F, Extended Data Fig. S11B, Table S13). These distal regions lay adjacent to immediate-early genes (*Egr1*, *Egr2*, *Egr3*), neuronal specification genes (*Ascl1*, *Bcl11a*, *Nrg3*) and epigenetic regulators (*Dnmt3a*, *Setd5*) (Fig. 5G). The stimulus-associated chromatin response within the pyramidal lineage is therefore graded by maturation state, with the larger response in newly formed neurons; whether this reflects a greater regulatory response per cell or the expanded representation of that population cannot be separated in these data.

### TE-derived stimulus-responsive regions are occupied by neurogenic transcription factors and lie near neuronal genes

Finally, we asked whether the TE-derived component of the response is preferentially associated with newly formed neurons and whether these sequences carry evidence of transcription-factor occupancy. Newly formed pyramidal neurons contained more TE-overlapping DARs (TE-DARs) than mature neurons at both promoter-proximal and distal positions: 1,934 versus 1,253, with 844 of the former located within +/-10 kb of a gene promoter (Fig. 6A, Extended Data Fig. S11C-D, Table S13).

**Figure 6.**
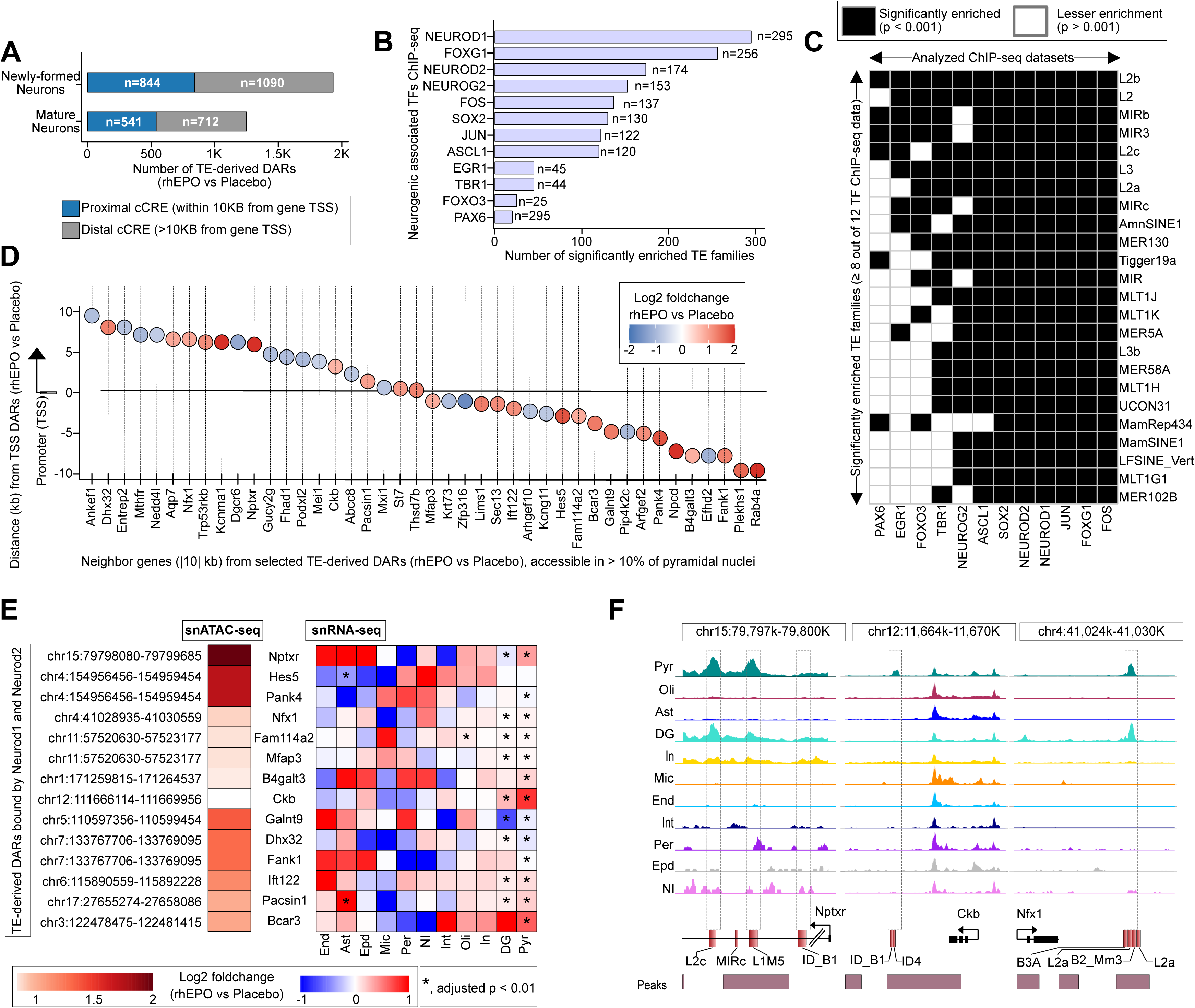
TE-derived stimulus-responsive regions are bound by neurogenic TFs and lie near differentiation and synapse genes. **(A)** Numbers of TE-derived DARs (rhEPO versus placebo) in newly formed and mature neurons, separated into proximal candidate cCREs (within 10 kb of a gene TSS, blue) and distal cCREs (>10 kb from a gene TSS, grey). Newly formed neurons harbour more TE-derived DARs than mature neurons. **(B)** Bar chart ranking neurogenic TFs by the number of TE families significantly enriched within their ChIP-seq binding sites. Each bar represents the total count of enriched TE families associated with the indicated TF. **(C)** Matrix of significantly enriched TE families across TF ChIP-seq datasets. Black squares denote significant enrichment (p < 0.001), white squares no significant enrichment (p > 0.001). TE families including L2b, L2, MIRb and MIR3 are associated with multiple neurogenic TFs. **(D)** Neighbouring genes within +/-10 kb of selected TE-derived DARs (rhEPO versus placebo) accessible in >10% of pyramidal nuclei. Each circle represents the closest DAR relative to the TSS, with position on the y-axis showing distance (kb) from the TSS and colour indicating log2 fold change (red = increased, blue = decreased accessibility under rhEPO). **(E)** Concordance between TE-derived DARs bound by neurogenic TFs and their nearest genes. Left: genomic coordinates of DARs identified by snATAC-seq, with colour scale indicating log2 fold change (rhEPO versus placebo). Right: heatmaps of matched gene expression from snRNA-seq across hippocampal cell types. Asterisks denote significant transcriptional changes (FDR < 0.01). **(F)** Genome browser snapshots showing TE-derived DARs near neurogenic genes. Representative loci include Nptxr, Ckb and Nfx1. Tracks display chromatin accessibility across major hippocampal lineages, with dashed boxes marking TE-overlapping elements that show lineage-restricted accessibility. Genes are shown in black and TEs highlighted in red.

To assess the regulatory context of these sites, we intersected TE-DARs with published ChIP-seq profiles for 12 neurogenic transcription factors (FOXG1, PAX6, ASCL1, NEUROG2, NEUROD1, NEUROD2, SOX2, TBR1, EGR1, FOXO3, FOS and JUN). Using established permutation tests^55^, hundreds of TE-DARs were bound by one or more of these factors, exceeding chance expectation (Fig. 6B, Fig. 7A-C, Extended Data Fig. S12, Table S14). NEUROD1 and FOXG1 showed the most extensive binding across TE families (295 and 256 families, respectively; p < 0.001). A core set of 24 TE families was enriched across multiple neurogenic factors, comprising LINEs (L2, L3), LTR retrotransposons (MER130 and MLT1 series) and SINE elements (AmnSINE1, MamSINE1) (Fig. 6C, Extended Data Fig. S12-S13). Several of these families have documented enhancer activity in the brain: MER130 elements harbour consensus NEUROD/NEUROG binding motifs and function as enhancers during mouse neocortical development^56^, and AmnSINE1 has well-documented enhancer activity in mammalian brain development. The present data show that TE families with prior enhancer annotation are also over-represented among stimulus-responsive accessible regions in the adult hippocampus, and that they carry binding for the corresponding factors in independent datasets.

**Figure 7.**
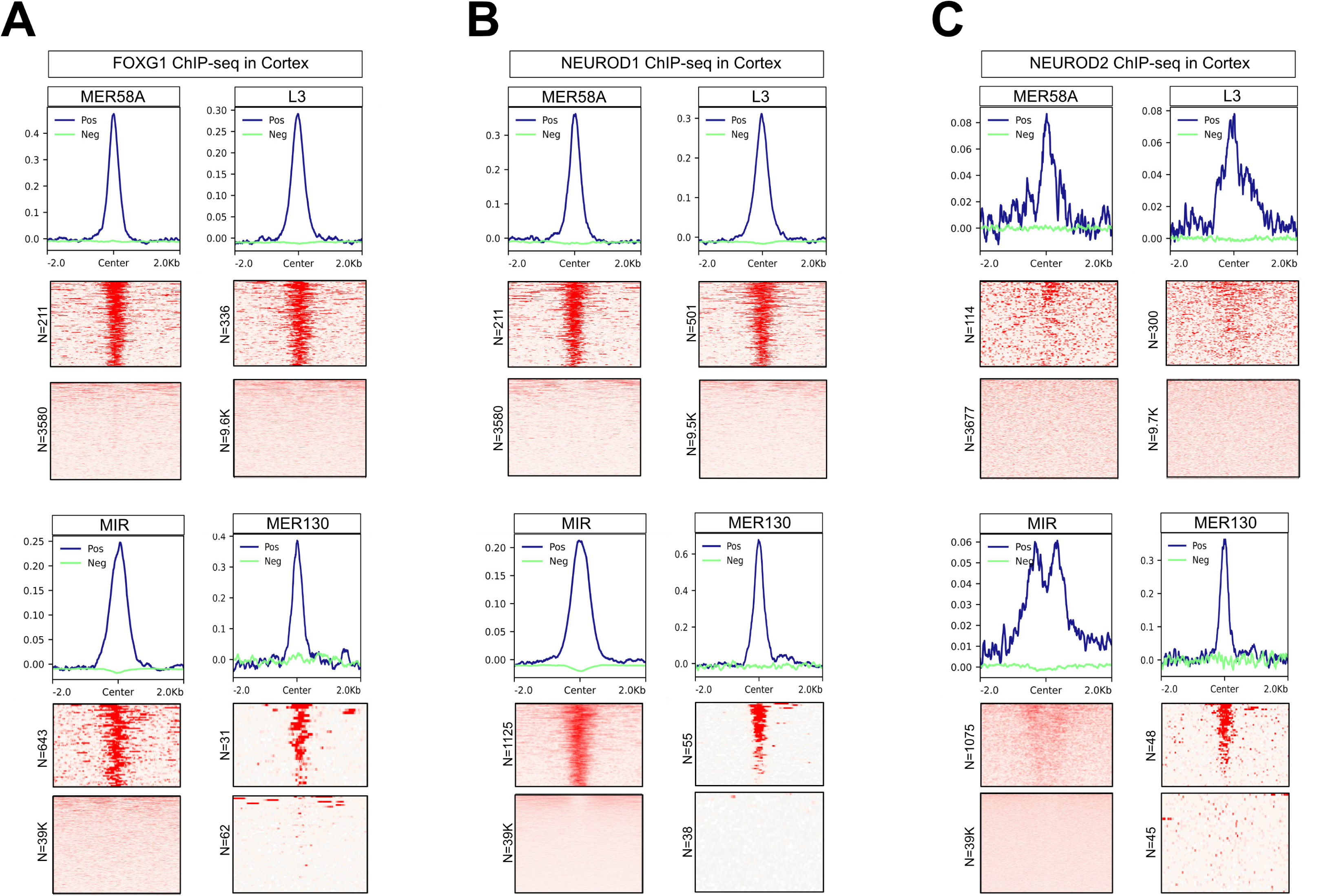
FOXG1, NEUROD1 and NEUROD2 ChIP-seq signal is enriched over defined TE subfamilies. **(A)** FOXG1 ChIP-seq aggregate signal profiles (top) and corresponding heatmaps (bottom) centred on representative repeat element families (MER58A, L3, MIR, MER130). “Positive” (Pos, blue) represents transposable elements overlapping FOXG1 ChIP-seq peaks, whereas “Negative” (Neg, green) indicates transposable elements without peaks. Each heatmap displays signal density within +/-2 kb of the element centre. **(B)** NEUROD1 ChIP-seq aggregate signal profiles and heatmaps across the same TE families (MER58A, L3, MIR, MER130). **(C)** NEUROD2 ChIP-seq meta-profiles and heatmaps over the same TE families. Together, these analyses show that specific TE subfamilies carry neurogenic TF occupancy, identifying them as candidate cis-regulatory modules of the neurogenic transcriptional program.

To relate TE sites to genes, we examined the chromatin state of these loci in their vicinity. Filtering for regions accessible in >10% of cells in at least one condition, 42 TE-containing DARs in the rhEPO versus placebo comparison lay within 10 kb of an annotated TSS (Fig. 6D, Table S15). Gene ontology analysis of the nearest genes to TF-bound TE-DARs returned enrichment for biological processes related to neurogenesis and synaptic integration (Extended Data Fig. S13, Table S16), and the nearest genes included *Hes5*, *Kcnma1*, *Pacsin1*, *Ankef1*, *Rab4a*, *Nptxr* and *Nedd4l*, which have documented roles in neuronal differentiation, maturation and synapse development (Fig. 6D-E). Accessibility at TF-bound TE-DARs was increased in rhEPO samples (FDR < 0.01), with concordant transcriptional upregulation of neighbouring genes in newly formed neurons and dentate gyrus (Fig. 6E). The candidate loci listed above were accessible specifically in rhEPO newly formed neurons, and the corresponding genes were upregulated in the matched snRNA-seq data (Fig. 6E-F).

Taken together, TE-derived sequences from defined families are over-represented among stimulus-responsive accessible regions in newly formed pyramidal neurons, are bound by neurogenic transcription factors in independent ChIP-seq data above chance expectation, and lie near genes whose expression changes in the same direction and in the same cells. These properties define TE-derived sequences as cCREs of the stimulated neurogenic program. Establishing that any of these elements is required for the expression of a neighbouring gene will require targeted silencing or deletion, which the present study does not provide.

## DISCUSSION

We have assembled a lineage-resolved atlas of chromatin accessibility in the adult mouse hippocampus and used it to describe the regulatory state of the tissue under a defined pro-neurogenic stimulus. Three features of the response are consistent in our analyses. First, chromatin remodelling is unevenly distributed across hippocampal lineages: more than half of all differentially accessible regions arise in pyramidal neurons, while dentate gyrus neurons and interneurons show a net loss of accessibility. Second, within the pyramidal lineage the response is graded by maturation state, with the larger set of differentially accessible regions in newly formed rather than mature neurons, and with a substantial distal component that transcript measurements do not report. Third, about 50% of the regions that gain accessibility overlap transposable elements, a proportion above the ∼20% baseline overlap we measure in the unperturbed hippocampal cCRE repertoire, and a defined subset of TE families within these regions is bound by neurogenic transcription factors in independent ChIP-seq datasets. We interpret these TE-derived regions as cCREs of a stimulus-responsive neurogenic programme. Throughout, rhEPO serves as an experimentally controlled entry point to that programme rather than as the object of study: its cellular consequences in this paradigm were established previously by single-nucleus transcriptomics^1,6^, and the contribution here is the regulatory layer underlying them.

### Chromatin accessibility reveals regulatory changes beyond gene expression

The chromatin profiles expose a substantial distal response that transcript measurements cannot capture. In newly formed pyramidal neurons, promoter-proximal regions represented a minority of the rhEPO-associated changes. Approximately 4,000 DARs occurred within ±10 kb of an annotated TSS, including regions near the immediate-early genes Egr1, Egr2, and Egr3; neuronal specification genes Ascl1, Bcl11a, and Nrg3; and epigenetic regulators Dnmt3a and Setd5. These distal elements constitute one of the largest single category of stimulus-responsive regions we detect, and none of them is reported in transcript data.

The atlas also identifies hippocampal cCREs missing from previous brain-wide annotations. Approximately 10.8% of hippocampal peaks were absent from a recent single-cell accessibility atlas of the adult mouse brain, and the *Slc1a2* locus illustrates the resolution at which the resource operates, with a promoter accessible across lineages and distal elements open only in astrocytes. Thus, the dataset distinguishes broadly accessible promoters from lineage-restricted distal elements that accompany cell-type-specific expression.

Chromatin and transcription changed coherently in the matched cell populations. DARs proximal to DEGs showed concordant directions of change, and the highlighted loci in newly formed neurons were accompanied by increased expression of neighbouring genes. Both modalities were measured at the same post-treatment time point. Their concordance, therefore, connects chromatin accessibility with transcription in the same cell types but does not resolve which change occurs first.

### rhEPO remodels chromatin preferentially in newly formed pyramidal neurons

The directional split between lineages is quite a robust feature of the dataset. Pyramidal neurons carried the dominant chromatin response to rhEPO, accompanied by smaller accessibility gains in oligodendrocytes, astrocytes, and microglia. Dentate gyrus granule neurons and interneurons instead showed a net loss of accessibility. In pyramidal neurons, this is accompanied by increased accessibility at promoters of neurogenic regulators, including *Ascl1* and *Neurog2*, together with less-characterized regulators such as *Zbtb16*, *Zbtb18*, *Fabp5*, *Mdga2* and *Dusp6*. We did not detect evidence of aberrant cell-cycle activation in non-neuronal lineages. This observation supports our previous conclusion that rhEPO shifts the fate and maturation of pre-existing precursors without inducing widespread cell division.

However, this interpretation of EPO-mediated cell-fate determination confers an obvious limitation. The reduced accessibility in interneurons parallels the previously reported reduction in their relative abundance and inhibitory potential. More generally, a change in accessibility measured across a lineage cannot be separated from a change in the composition of that lineage. The same ambiguity applies to the pyramidal response, where an expanded immature population and a larger per-cell response would produce similar aggregate signals.

This distinction is important when interpreting the apparent expansion of newly formed pyramidal neurons. These subpopulations were up to 5-fold more abundant in the chromatin dataset, whereas snRNA-seq under the same regimen showed a ∼2-fold (∼200%) enrichment of immature pyramidal neurons and prolonged treatment increased mature CA1 pyramidal neurons by ∼20%.

We therefore treat the ∼200% transcriptomic value as the reference estimate and the chromatin-derived value as an upper bound. Several non-exclusive explanations are available: chromatin profiling may detect immature neurons more sensitively than transcript-based assignment, if accessibility at neurogenic loci is established before the corresponding transcripts accumulate; nuclear recovery from immature and mature neurons may differ; and dissection or regional sampling may vary between samples, particularly as each snATAC-seq library was generated from pooled left hippocampi. Distinguishing these possibilities requires orthogonal quantification, such as histological counting of immature markers in the same animals, which we did not perform. The lineage selectivity of the chromatin response does not depend on the absolute magnitude of this expansion, and we build no argument on the 5-fold value.

### TE-derived regions contribute to the inducible neurogenic repertoire

TE-derived sequences were disproportionately represented among chromatin regions gaining accessibility under rhEPO. Approximately one-fifth of accessible regions overlapped TEs across the hippocampal cCRE repertoire, close to the ∼25% reported for mammalian cCREs^39^ and consistent with epigenomic surveys of brain regulatory elements^18,20^. This fraction rose to approximately 50% among regions gaining accessibility upon rhEPO treatment. In the aggregate comparison, these are predominantly rodent-specific SINEs of the B1/Alu and B2 families, including *ID_B1*, *B3* and *B1_Mus1/2*. In the lineage-specific analysis, enriched families include LTR retrotransposons of the ORR1E and MTD groups, the L2/L2a LINEs, the rodent B3 and B4/B4A SINEs, and MIR/MIRb. The analysis of independent ChIP-seq datasets narrows this to a core set of 24 families bound above chance by multiple neurogenic factors, with NEUROD1 and FOXG1 showing the broadest occupancy. This set comprises the L2 and L3 LINEs, the LTR33, MER130 and MLT1 elements, and the AmnSINE1 and MamSINE1 SINEs. Several of these families have documented enhancer activity in the brain. For instance, MER130 elements contain consensus NEUROD/NEUROG motifs and act as enhancers during mouse neocortical development, whereas AmnSINE1 elements function as enhancers during mammalian brain development.

Multiple lines of evidence nominate these TE-derived regions as cCREs. They are accessible in a stimulus-dependent and cell-type-restricted manner, they are occupied by the relevant transcription factors in independent datasets, they lie within 10 kb of genes with described roles in neuronal differentiation and synapse formation, including *Hes5*, *Kcnma1*, *Pacsin1*, *Ankef1*, *Rab4a*, *Nptxr* and *Nedd4l*, and those genes change expression in the same direction in the same cells. These observations support a model in which TE-derived sequences carrying neurogenic TF motifs become accessible during stimulation and contribute to the induced transcriptional program. This model remains to be tested by perturbing individual elements and measuring their effects on the proposed target genes.

The findings add an inducible adult-brain context to the broader observation that cell-type-specific transcription factors frequently occupy transposon-derived sequences. Previous work has largely emphasised TE exaptation during development and evolution^22,23,26,40^. Here, the same principle emerges in newly formed neurons responding to an external pro-neurogenic signal.

### Neurogenic regulons connect accessibility with neuronal maturation

Our integrative multi-omic analysis sheds light on the gene regulatory networks fine-tuned under rhEPO and how they intersect with TEs. Integration of the chromatin and transcriptomic profiles identified approximately a dozen regulons enriched under rhEPO against a largely shared regulatory background. These networks were driven by factors controlling neuronal differentiation and maturation, including NEUROD1, NEUROD2, FOXG1, PAX6, and JUN. Their motifs were enriched in regions gaining accessibility, some of which were TE-derived, while the corresponding transcripts increased in pyramidal lineages^1^. These relationships suggest a feed-forward organisation in which increased expression of neurogenic TFs accompanies greater accessibility of their binding sites. Because Pando infers regulons from co-accessibility and co-expression, this organisation represents a regulatory model, not a demonstrated circuit.

Maturation state further shaped this response. Newly formed pyramidal neurons contained more DARs than mature neurons (Table S13), although both populations showed similar proportions of accessibility gains. Regions distinguishing the two states were enriched near genes controlling neurogenesis and synapse formation. This pattern suggests that regulatory responsiveness narrows as pyramidal neurons mature, potentially because differentiation restricts the repertoire of elements available to external stimuli. The simultaneous expansion of the immature population prevents a complete separation of per-cell responsiveness from population abundance.

### EPO-responsive chromatin provides a basis for comparing neurogenic stimuli

Endogenous brain EPO increases under hypoxic and injury conditions and has been implicated in neuroprotection^11^. Adult hippocampal neurogenesis, first recognized in rodents in the 1960s^57,58^ and three decades later was shown in humans^59^, and recently the proliferative neuro progenitor cells are identified in adult hippocampus^3^. Clinical and preclinical studies further show that rhEPO can improve learning and memory independently of erythropoiesis^7,9,60^. The chromatin states described here raise the testable possibility that hypoxia, injury, or other pro-neurogenic stimuli engage overlapping regulatory elements. This study did not examine hypoxia, HIF-dependent signalling, or metabolic state, and the overlap between these responses remains unknown.

For the same reason, we frame the translational implication narrowly. The cell-type-resolved catalogue of accessible regions can now support comparisons across neurogenic stimuli and prioritize loci for functional testing. Its annotations connect each region to genomic context, TE origin, motif content, and proximal genes. In particular, the elements selectively accessible in newly formed neurons provide candidates for testing whether induced chromatin states can be manipulated to influence neuronal maturation. Such applications remain prospective because this study neither establishes the function of an individual element nor examines a disease model.

### Limitations of the study

Our study captures one time point after completion of the treatment and therefore cannot resolve the temporal ordering of chromatin and transcriptional changes, or distinguish transient from sustained accessibility changes. Chromatin accessibility reports regulatory potential but does not measure histone modifications, DNA methylation, or three-dimensional chromatin contacts. We also did not perturb individual regions. Functional validation will require approaches such as CRISPR-mediated deletion of selected TE-derived elements followed by measurement of the proposed target genes. Similarly, the target genes are assigned by proximity rather than by measured physical contact. The design comprises four libraries per condition, each generated from pooled left hippocampi, with the matched snRNA-seq data derived from pooled right hippocampi (GSE220522); pooling limits our ability to model between-animal variability. Finally, the study uses one species, male mice of a single age range, and one dosing regimen, so the generality of the response across stimuli, sexes, regions and species remains to be tested.

## MATERIALS AND METHODS

### Ethical approval

All mouse experiments were approved by the local Animal Care and Use Committee (Niedersächsisches Landesamt für Verbraucherschutz und Lebensmittelsicherheit, LAVES - AZ 33.19-42502-04-17/2393) and conducted in strict accordance with the German Animal Protection Law. Every effort was made to minimize the number of mice used and their suffering.

### Experimental model and treatments

Male C57BL/6N mice received 11 intraperitoneal injections of recombinant human erythropoietin (rhEPO, 5000 IU/kg body weight) or placebo (0.01 mL/g of solvent) every other day for 3 weeks, starting at postnatal day 28 (P28). In total, 23 male mice were used (rhEPO = 11, placebo = 12). Twenty-four hours after the final injection (P49), all animals were sacrificed by cervical dislocation. For each sample, 2 right hippocampi from mice of the same treatment group were pooled into a single tube (except for one rhEPO sample, which included only one hippocampus), resulting in 6 biological replicates per condition (N = 6 rhEPO, N = 6 placebo) for snRNA-seq (GSE220522). Likewise, the left hippocampi from 4 tubes of each group (N = 4 rhEPO, N = 4 placebo; tube labels A1, A2, A4, A6, B7, B8, B9, B11) were used for snATAC-seq. snATAC-seq library construction was performed for these 8 hippocampal samples using Chromium Single Cell ATAC Library and Gel Bead Kit v2 chemistry.

### Single-cell ATAC-seq library processing and alignment

snATAC-seq libraries were generated and processed using the Cell Ranger ATAC pipeline (v2.1.0, 10x Genomics). Raw reads from 8 hippocampal samples were aligned to the mm10 reference genome (refdata-cellranger-arc-mm10-2020-A-2.0.0). Peak-barcode matrices and fragment files were generated using cellranger-atac count and used for downstream analysis.

### Unified-peak set workflow

Downstream analysis was performed in R using the Signac (v1.13.0) and Seurat (v5.1.0) packages. Individual peak files were merged into a unified peak set using reduce() from the GenomicRanges package, excluding peaks <20 bp or >10,000 bp. Cells with fewer than 500 fragments in peaks were filtered out. Chromatin assays were constructed for each sample using FeatureMatrix().

Each Seurat object was annotated using EnsDb.Mmusculus.v79, and quality control (QC) metrics were computed: transcription start site (TSS) enrichment, nucleosome signal, percent reads in peaks, and blacklist ratio (based on blacklist_mm10). Cells were filtered using quantile-based thresholds, excluding outliers (below the 2nd or above the 98th percentile for key metrics) to remove low-quality cells and potential multiplets.

Filtered objects were merged, then the merged object was processed with normalization using RunTFIDF(), dimensionality reduction via latent semantic indexing (RunSVD()), batch correction with Harmony (RunHarmony() using LSI dims 2-30), and UMAP embedding. Clusters were identified with FindNeighbors() and FindClusters() with resolution 0.8 and algorithm 1. Gene activity scores were calculated via GeneActivity() and log-normalized with NormalizeData() (scale factor = median of nCount_Activity). A preprocessed single-nucleus RNA-seq dataset^1^ served as a reference for cell-type annotation. Unmatching samples (A3, A5, B10, B12) were excluded. Anchors were computed using canonical correlation analysis (CCA) to predict cells in the snATAC-seq dataset.

### Sample-specific peak set workflow

In addition to the unified peak set approach, we conducted a complementary analysis using sample-specific peak sets generated by Cell Ranger ATAC. For each sample, peak-barcode matrices and fragment files were independently loaded, and chromatin assays were constructed using a minimum cell threshold of min.cells = 10. Seurat objects were generated separately for each sample and subsequently merged for integrated downstream analysis. Gene annotations were added using the EnsDb.Mmusculus.v79 database. QC metrics were calculated for each cell, including TSS enrichment, nucleosome signal, percent reads in peaks, and the blacklist ratio. To ensure high-quality samples, cells were filtered using the following criteria: nCount_peaks between 3,000 and 30,000, pct_reads_in_peaks >15%, blacklist_ratio <0.05, nucleosome_signal <4, and TSS.enrichment >3.

Dimensionality reduction and clustering were performed: normalization with RunTFIDF(), feature selection via FindTopFeatures(min.cutoff = ‘q0’), latent semantic indexing (LSI) using RunSVD() on dimensions 2 through 8, and UMAP embedding via RunUMAP(). Clusters were identified using FindNeighbors() and FindClusters() with algorithm 1. Gene activity scores were computed using GeneActivity() and log-normalized with NormalizeData(). Cell-type annotations previously inferred from the unified peak set analysis were transferred to the sample-specific peak set dataset by matching shared cell barcodes. Cells without annotation were excluded from further analysis.

For analyses related to the pyramidal cluster, pyramidal neurons were subset from the unified peak set dataset and subjected to reclustering to refine subtype resolution. To facilitate annotation, these cells were integrated with a previously annotated single-nucleus RNA-seq reference dataset^1^ using CCA, with 30 dimensions and 10 nearest neighbours employed to identify cross-modality anchors. Uncharacterized or ambiguous Seurat clusters were either removed or reannotated based on their transcriptomic profiles and proximity to well-defined clusters in the reference dataset. The finalized annotations were then transferred to the corresponding pyramidal neuron populations within the sample-specific peak set dataset by matching shared cell barcodes.

### Differential accessibility analysis

All rhEPO- and placebo-derived cells were combined within each cluster (Seurat cluster, higher-level cell-type annotation, or pyramidal subcluster). We then executed FindMarkers (test.use = “LR”, min.pct = 0.01, latent.vars = “nCount_peaks”) in a one-versus-rest fashion to identify peaks that are intrinsically enriched or depleted in a given cluster relative to all other clusters. We used 0.01 as the min.pct threshold to capture accessibility shifts in smaller subfractions of a population. Peaks passing FDR < 0.01 were considered cell-type DA peaks. Furthermore, to pinpoint treatment-dependent changes, we stratified cells by condition inside each cluster and ran FindMarkers() with min.pct = 0.01 in the LR framework, with depth correction applied. Peaks with FDR < 0.01 were classified as condition DA peaks. For differences between defined clusters (cell-type DA peaks) and for treatment-dependent changes (condition DA peaks), summits (peak mid-points) were annotated via TxDb.Mmusculus.UCSC.mm10.knownGene to the closest gene’s promoter, binned by genomic context (promoter, exon, intron, intergenic) and intersected with RepeatMasker (mm10) to identify and quantify overlap with TEs. Low-complexity, satellite, RNA, rRNA, tRNA, scRNA, snRNA, RC, srpRNA and ambiguous repeats (entries flagged with “?”, such as “LINE?”) were excluded from all TE analyses unless explicitly stated.

### Cluster-resolved motif enrichment

Chromatin dynamics were analyzed in two complementary tiers: chromatin accessibility differences between defined clusters, and treatment-dependent changes. Vertebrate position-frequency matrices from the JASPAR-2020 CORE collection were used for motif analysis. For every Seurat cluster, higher-level cell-type annotation, or pyramidal subcluster, DA peaks identified using FindMarkers were filtered at a min.pct threshold of 0.10, positive only, and FDR < 0.05. Clusters contributing >= 10 significant peaks were subjected to motif enrichment with FindMotifs(). Likewise, within the same clustering framework, cells were split by condition, and DA peaks filtered with a min.pct threshold of 0.10, positive only, and FDR < 0.05 were used for motif analysis. For any cluster yielding >= 10 rhEPO-responsive peaks, FindMotifs() was run to identify transcription-factor motifs enriched in rhEPO-gained chromatin.

### Transcription-factor ChIP-seq integration

Genome-wide binding profiles for 12 neurodevelopmental transcription factors (ASCL1, EGR1, FOS, FOXG1, FOXO3, JUN, NEUROD1, NEUROD2, NEUROG2, PAX6, SOX2, TBR1) were downloaded from GEO (Table S17). Peak files were lifted over to the mm10 assembly where necessary (UCSC liftOver) and collapsed into non-redundant consensus regions with the bedtools merge function. To ask whether individual TFs preferentially occupy repeat-derived regulatory DNA, we quantified intersections between each consensus ChIP peak set and RepeatMasker annotations using regioneR; 1000 permutations provided empirical P values and observed/expected overlap ratios. TE-associated peaks were assigned to the nearest transcription start site with ChIPseeker (TxDb.Mmusculus.UCSC.mm10.knownGene) and subjected to GO biological-process enrichment with clusterProfiler::enrichGO (BH-adjusted q <= 0.01). Next, TF peaks were overlapped with the 11 broad cell-class DA peak lists and the refined pyramidal-lineage DA peaks identified in our snATAC-seq analysis.

## DATA AND CODE AVAILABILITY

All raw FASTQ files, fragment files, and count matrices underlying the present study have been deposited in the Gene Expression Omnibus and will be released upon publication. Accession numbers will also be listed at https://github.com/umutcakir/ATAC_EPO_vs_Placebo. Interactive visualizations of our scATAC-seq, snRNA-seq, and Seurat objects are available on the University of California, Santa Cruz (UCSC) Cell Browser under the accession name “tenet” (https://tenet.cells.ucsc.edu/). Intermediate files and supplementary data are available from the digital repository, Zenodo, DOI: 10.5281/zenodo.17635636. Code for data processing, quality control, clustering, differential-accessibility testing, motif analysis, and TF-ChIP overlap is available at https://github.com/umutcakir/ATAC_EPO_vs_Placebo.

## ACKNOWLEDGEMENTS AND FUNDING

This work was supported by the Centre National de la Recherche Scientifique (CNRS) through a Chaire de Professeur Junior (CPJ) awarded to M.S. for the project “Biologie des systèmes en physiopathologie.” and FRNEM welcome grant (FRN202509051056). H.E. was supported by the Deutsche Forschungsgemeinschaft (DFG, German Research Foundation) via the DFG Center for Nanoscale Microscopy and Molecular Physiology of the Brain (CNMPB). Research in the labs of HE and KAN is funded by SFB TRR 274/2 - 408885537 project C01 (to H.E. and K.A.N.). U.C. received support from the IMPRS-Genome Science PhD program. Research in the labs of D.G., R.K. and K.A.N. is supported by the Adelson Medical Research Foundation.

## AUTHOR CONTRIBUTIONS

Concept, design, supervision: MS, HE. Funding acquisition: MS, HE, KAN. Drafting manuscript: UC, MS, HE. Display items: UC, MS. Data acquisition and generation: UC, FM, SYA, UJB, VB, MS, RK, DG. Data analyses and interpretation: UC, RK, MS. All authors read and approved the final version of the manuscript.

## DECLARATION OF INTERESTS

The authors declare no competing financial or other interests in connection with this article.

## EXTENDED DATA FIGURE LEGENDS

**Extended Data Fig. S1.**
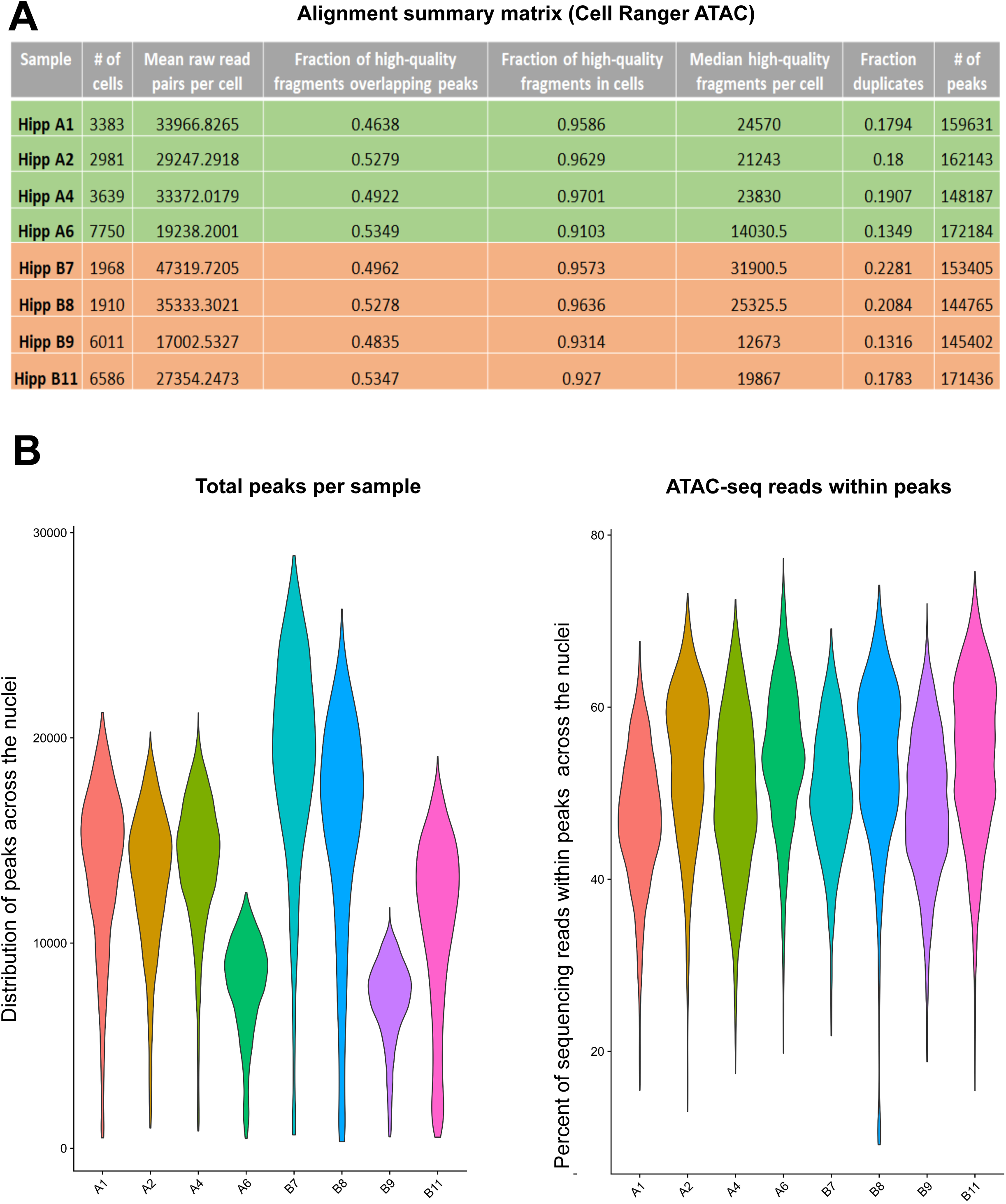
Quality control metrics derived from Cell Ranger ATAC output. **(A)** Summary of snATAC-seq quality metrics obtained from Cell Ranger ATAC output. For each sample the table shows the number of high-quality nuclei passing filtering, mean raw read pairs per nucleus, fraction of high-quality fragments overlapping called peaks, fraction of high-quality fragments confidently assigned to nuclei, median high-quality fragments per nucleus, duplicate fragment fraction, and total number of peaks detected. **(B)** Violin plots showing (left) the distribution of total peaks detected per nucleus and (right) the percentage of sequencing reads falling within peaks across nuclei for each sample.

**Extended Data Fig. S2.**
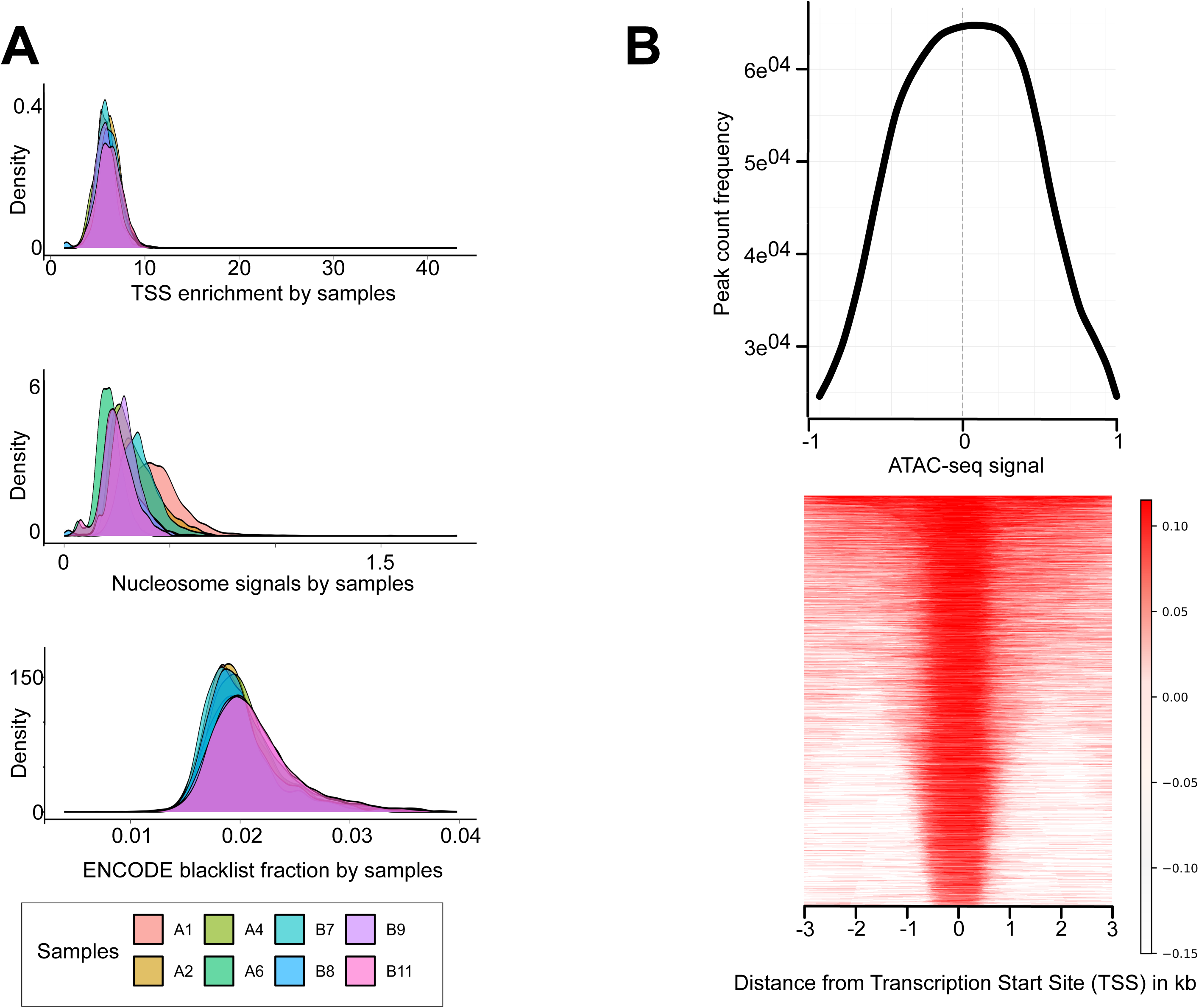
Additional quality control profiles for snATAC-seq libraries. **(A)** Density distributions of three core snATAC-seq quality metrics across samples: TSS enrichment scores, nucleosome signal estimates and ENCODE blacklist fractions. **(B)** Aggregate ATAC-seq signal profile centred on transcription start sites (top). The heatmap (bottom) displays ATAC-seq fragment density within +/-3 kb of TSSs for all peaks.

**Extended Data Fig. S3.**
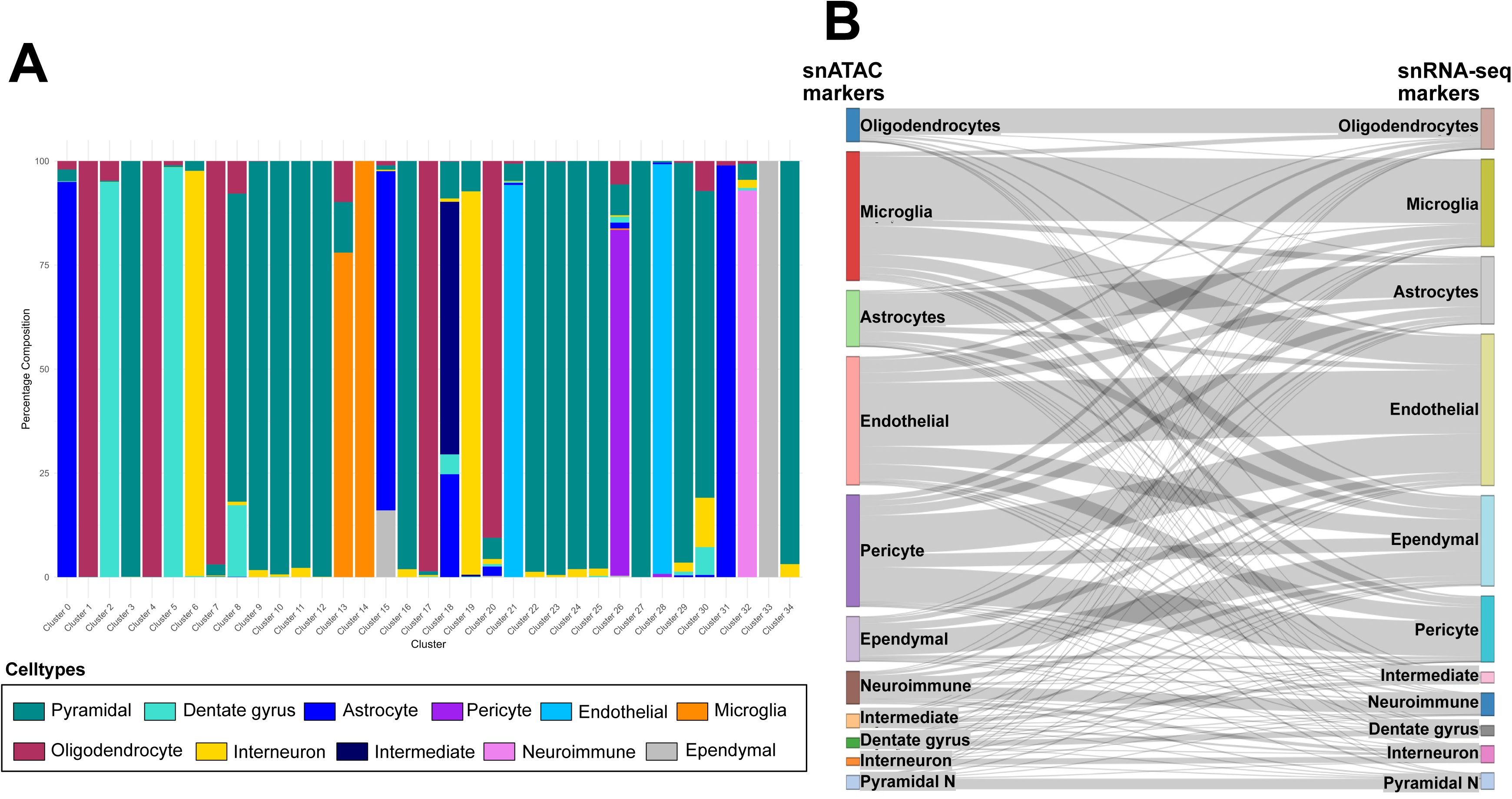
Cross-modality cell-type annotation and correspondence between snATAC-seq and snRNA-seq datasets. **(A)** Stacked bar plot showing the percentage composition of annotated cell types within each Seurat snATAC-seq cluster. Colours correspond to major hippocampal lineages. **(B)** Sankey diagram comparing cell-type annotations derived from snATAC-seq markers (left) with those obtained from snRNA-seq markers (right). Flow widths represent the degree of correspondence between modalities.

**Extended Data Fig. S4.**
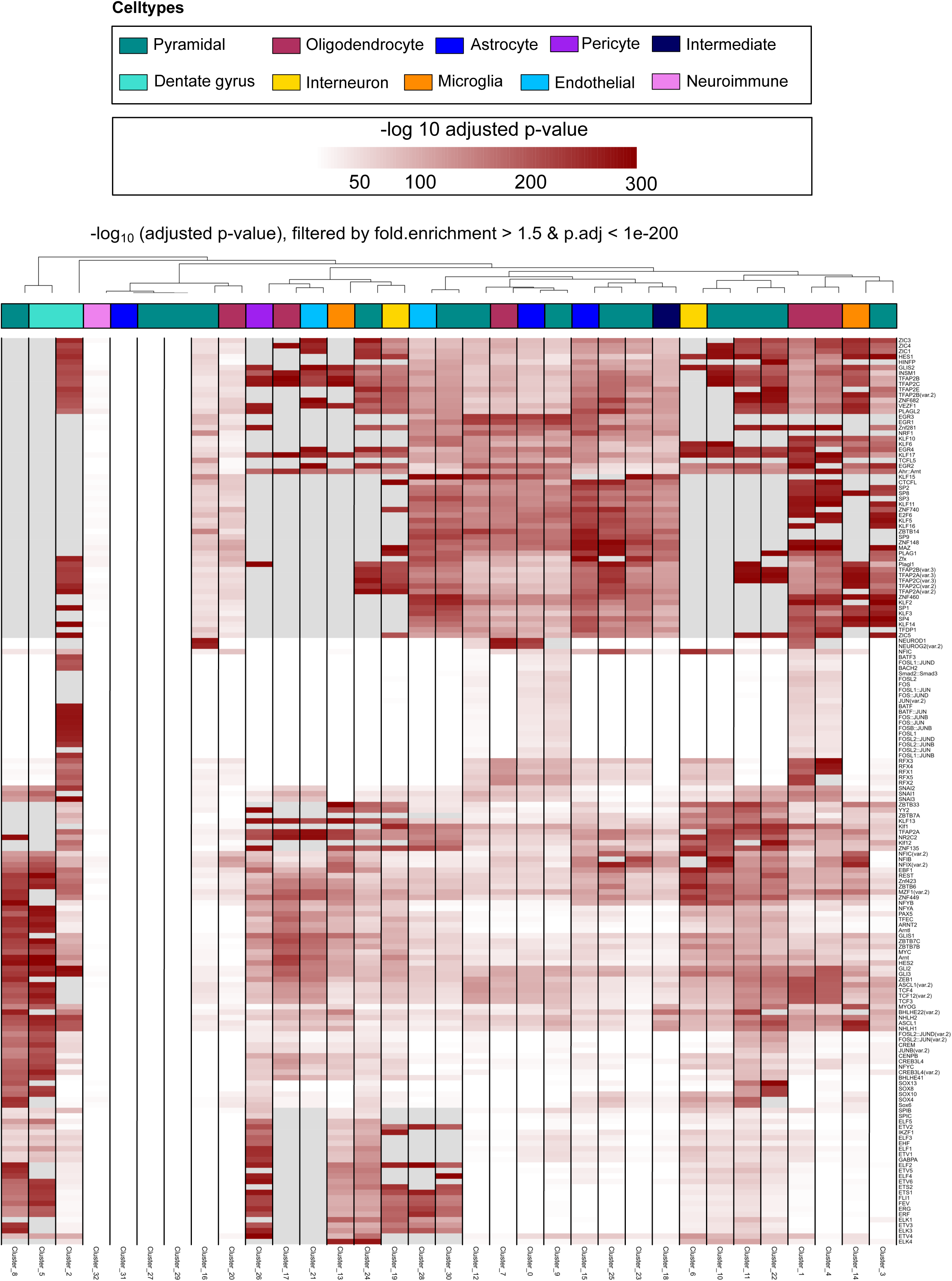
Transcription factor motif enrichment across Seurat snATAC-seq clusters. Heatmap showing enriched transcription factor motifs across Seurat-defined snATAC-seq clusters. Motifs were filtered for fold enrichment >1.5 and adjusted p-value <1 x 10-200. Each column represents an individual Seurat cluster annotated by its corresponding cell-type identity (top colour bar); each row shows a transcription factor motif, with colour intensity indicating the -log10 adjusted p-value of enrichment.

**Extended Data Fig. S5.**
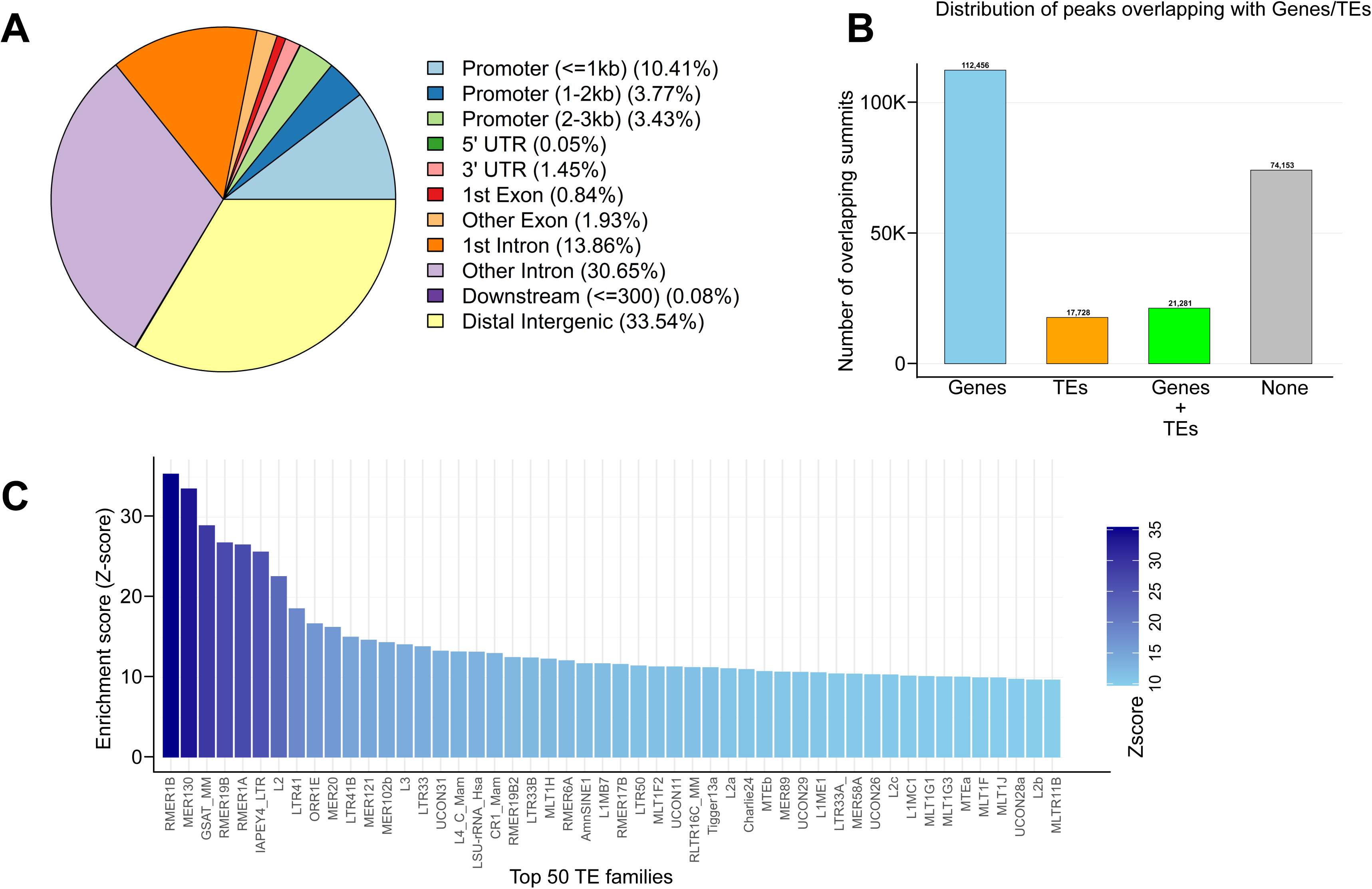
Genomic distribution of snATAC-seq peaks and enrichment of transposable element families. **(A)** Pie chart summarising the genomic annotation of all snATAC-seq peak summits. **(B)** Bar plot showing the number of peak summits overlapping annotated genes, annotated TEs, both, or neither. **(C)** Z-scores for the enrichment of the top 50 TE families overlapping snATAC-seq peak summits.

**Extended Data Fig. S6.**
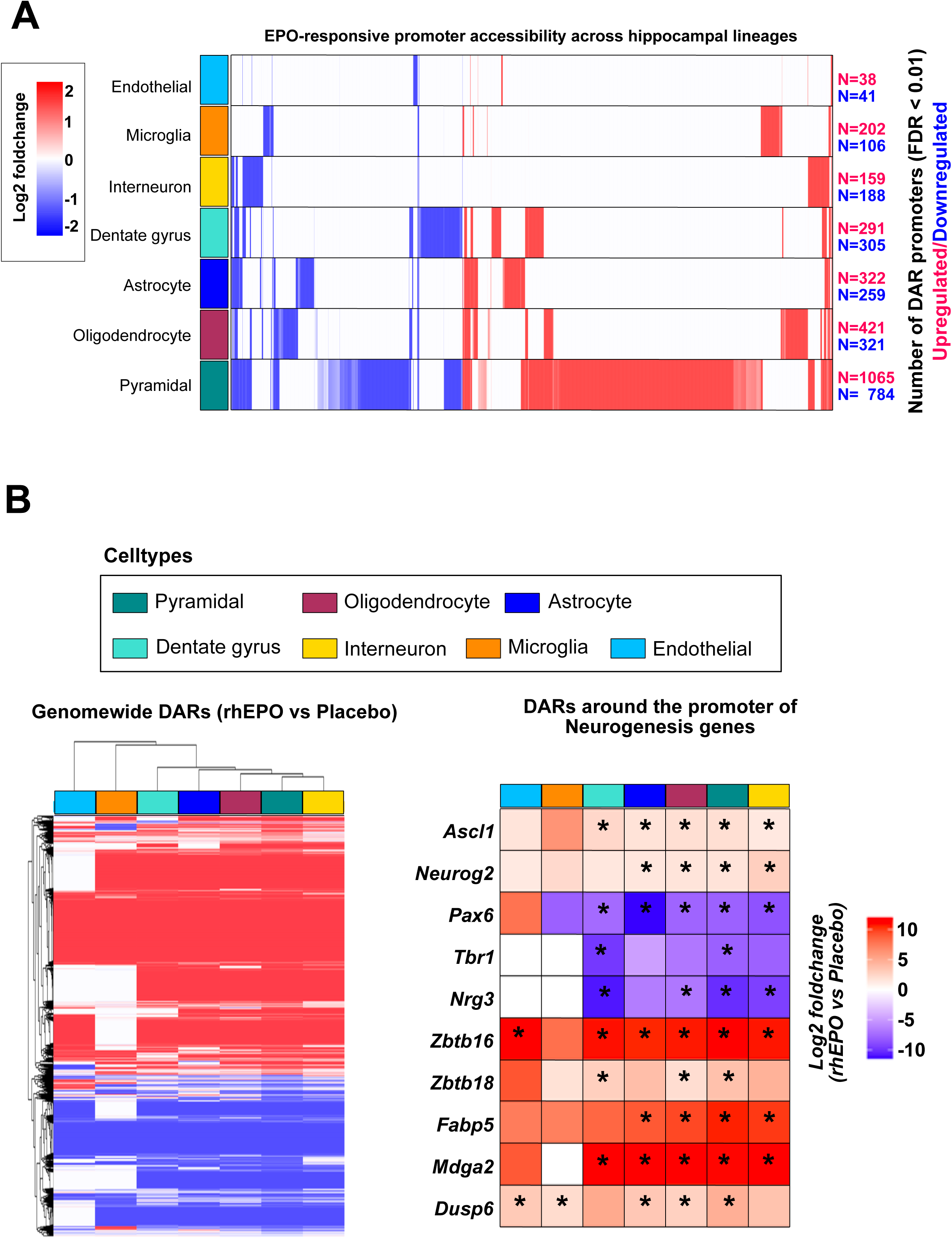
rhEPO-responsive promoter accessibility across hippocampal lineages. **(A)** Heatmap showing log2 fold change in promoter accessibility (FDR < 0.01) between rhEPO- and placebo-treated samples across major hippocampal cell types, with the total number of increased and decreased promoter DARs for each lineage shown at right. **(B)** Differential accessibility heatmaps summarising genome-wide DARs (left) and promoter-specific DARs for selected neurogenesis-related genes (right) across hippocampal cell types; genes shown are Ascl1, Neurog2, Pax6, Tbr1, Nrg3, Zbtb16, Zbtb18, Fabp5, Mdga2 and Dusp6. Asterisks denote significant change upon rhEPO treatment (FDR < 0.01).

**Extended Data Fig. S7.**
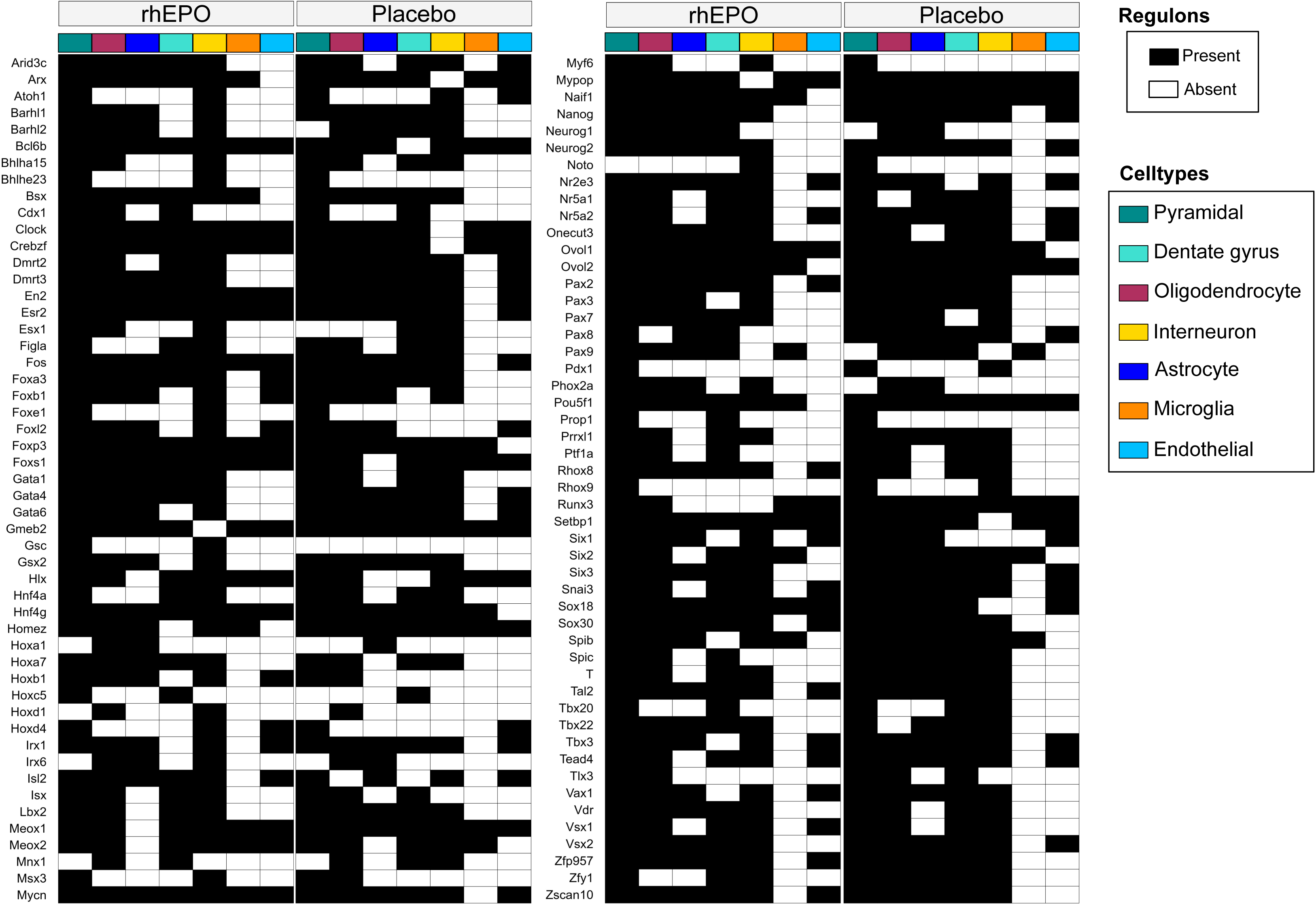
Regulon activity across hippocampal lineages under rhEPO and placebo conditions. Binary heatmap showing the presence or absence of Pando-inferred regulons across major hippocampal cell types for rhEPO- and placebo-treated conditions. Black boxes indicate regulons inferred as active in a given lineage, white boxes their absence.

**Extended Data Fig. S8.**
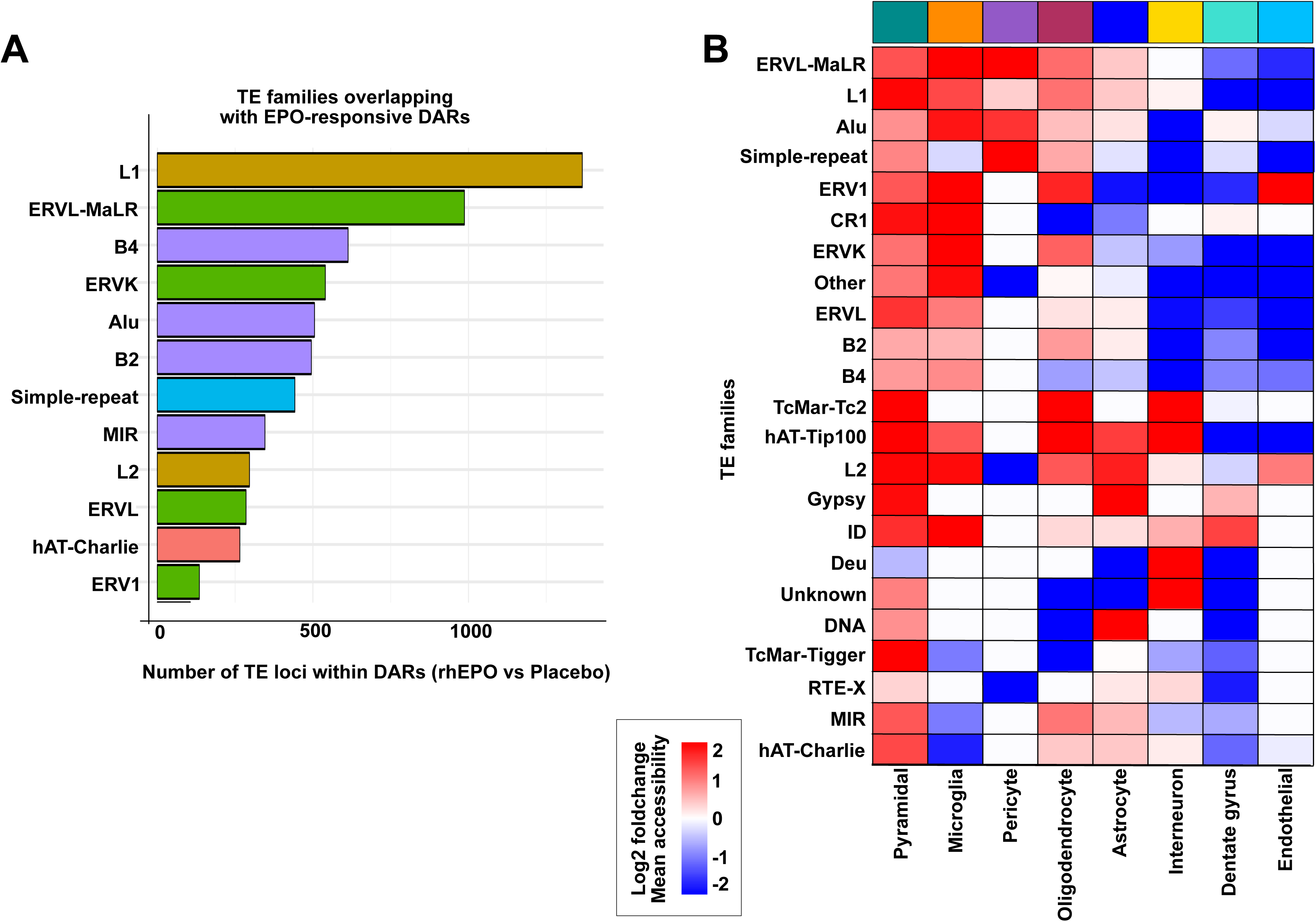
Transposable element families enriched within rhEPO-responsive chromatin regions. (**A)** Bar plot showing the number of TE loci overlapping DARs identified between rhEPO- and placebo-treated samples, ranked by the number of overlapping loci, with contributions from L1, ERVL-MaLR, B4, ERVK, Alu, B2 and other repeat classes. **(B)** Heatmap displaying the log2 fold change in accessibility for TE-associated DARs across major hippocampal cell types.

**Extended Data Fig. S9.**
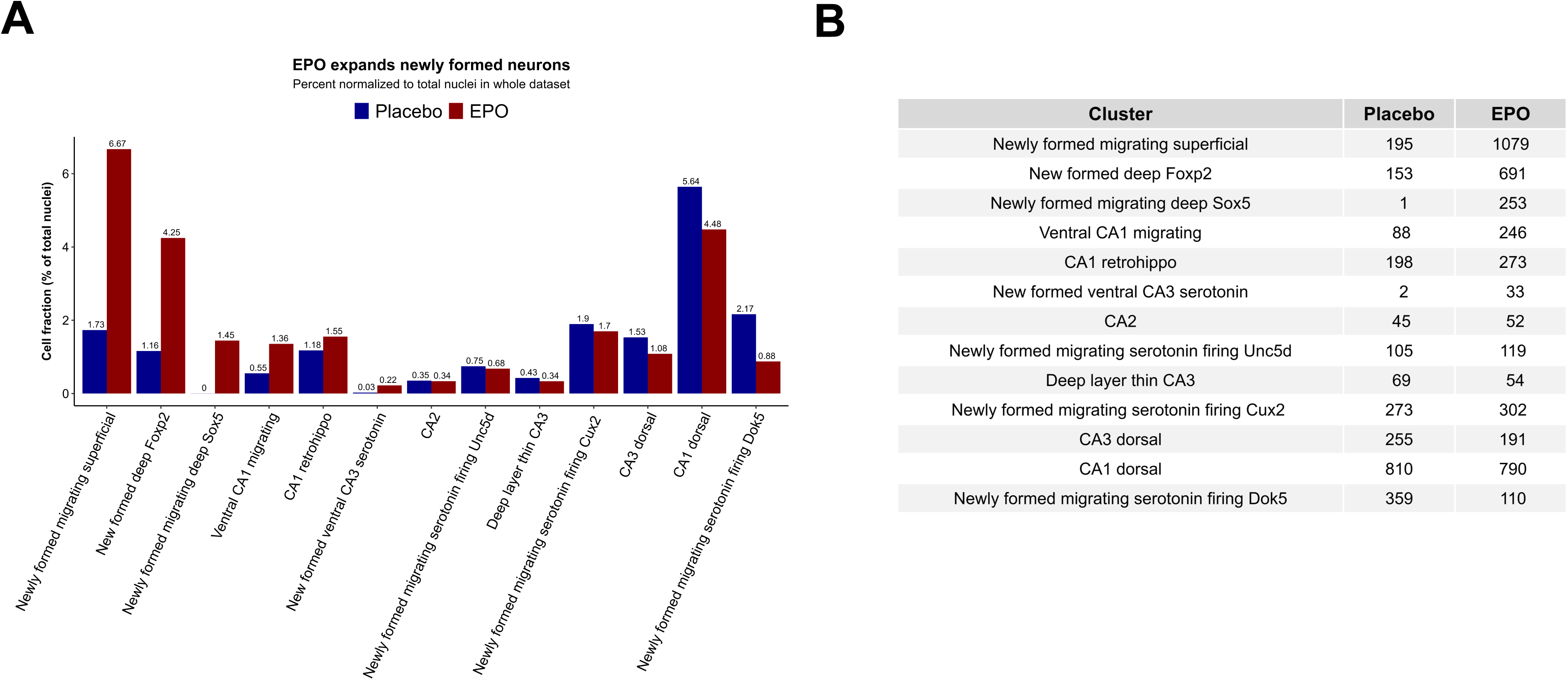
Abundance of newly formed pyramidal neuron populations under rhEPO treatment. **(A)** Bar plot showing the relative fraction of newly formed pyramidal neuron subtypes in placebo-and rhEPO-treated hippocampi, normalized to the total number of nuclei, across early-stage neuronal populations including superficial migrating, deep-layer Foxp2+ and migrating deep Sox5+ neurons, ventral CA1 migrating, CA1 retrohippocampal, and ventral CA3 serotonin neuron subtypes. **(B)** Table summarising the absolute number of nuclei assigned to each newly formed pyramidal neuron cluster under placebo and rhEPO conditions. These proportions are compositional estimates from recovered nuclei and are interpreted alongside the previously reported ∼200% transcriptomic enrichment^1^ (see Discussion).

**Extended Data Fig. S10.**
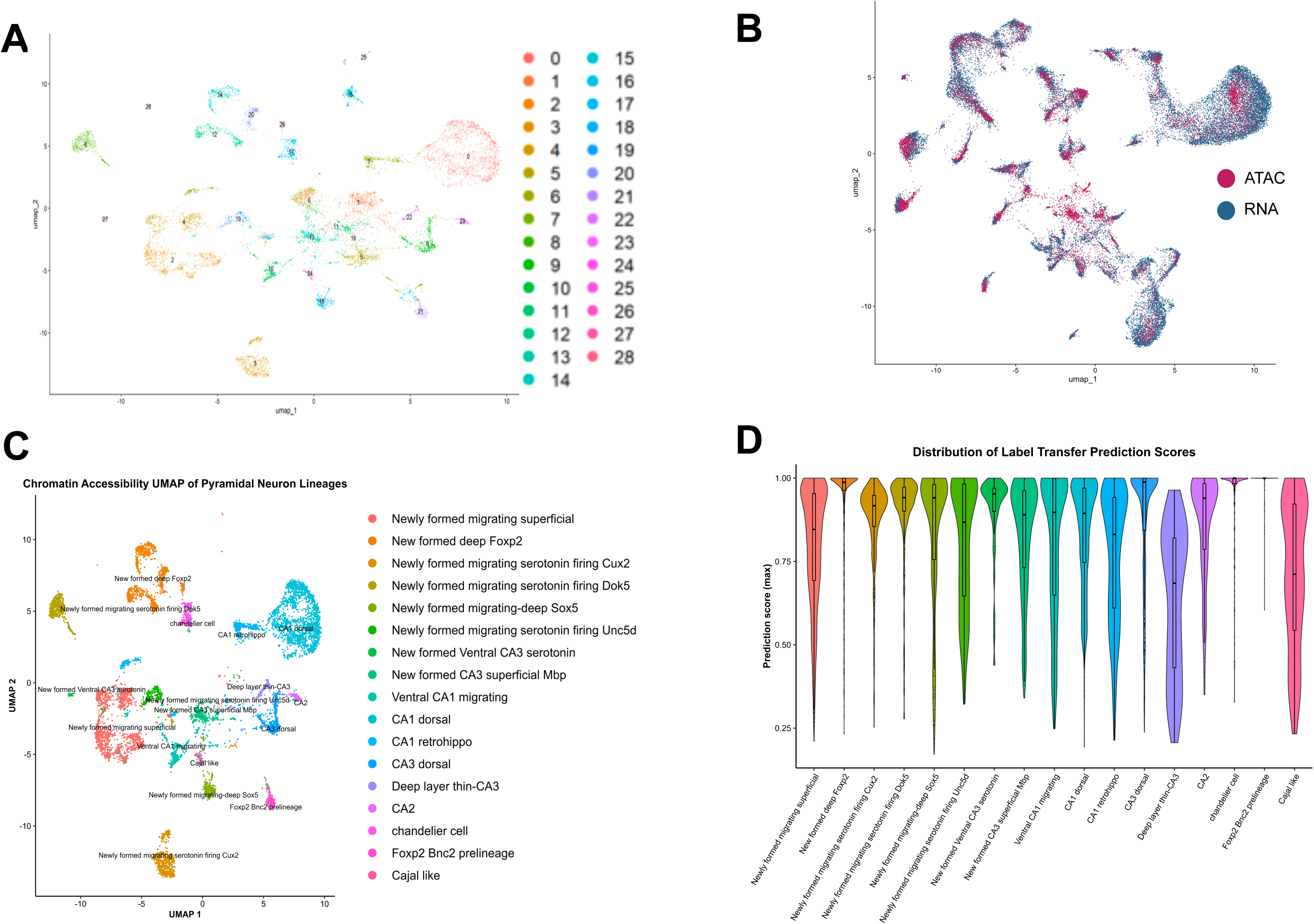
Integration of snATAC-seq and snRNA-seq datasets and annotation of pyramidal neuron lineages. **(A)** UMAP representation of Seurat-defined chromatin accessibility clusters from snATAC-seq. **(B)** Joint UMAP embedding of snATAC-seq (red) and snRNA-seq (blue) datasets. **(C)** UMAP visualization of chromatin accessibility profiles for annotated pyramidal neuron sublineages. **(D)** Violin plots of label transfer prediction scores for pyramidal neuron subtypes.

**Extended Data Fig. S11.**
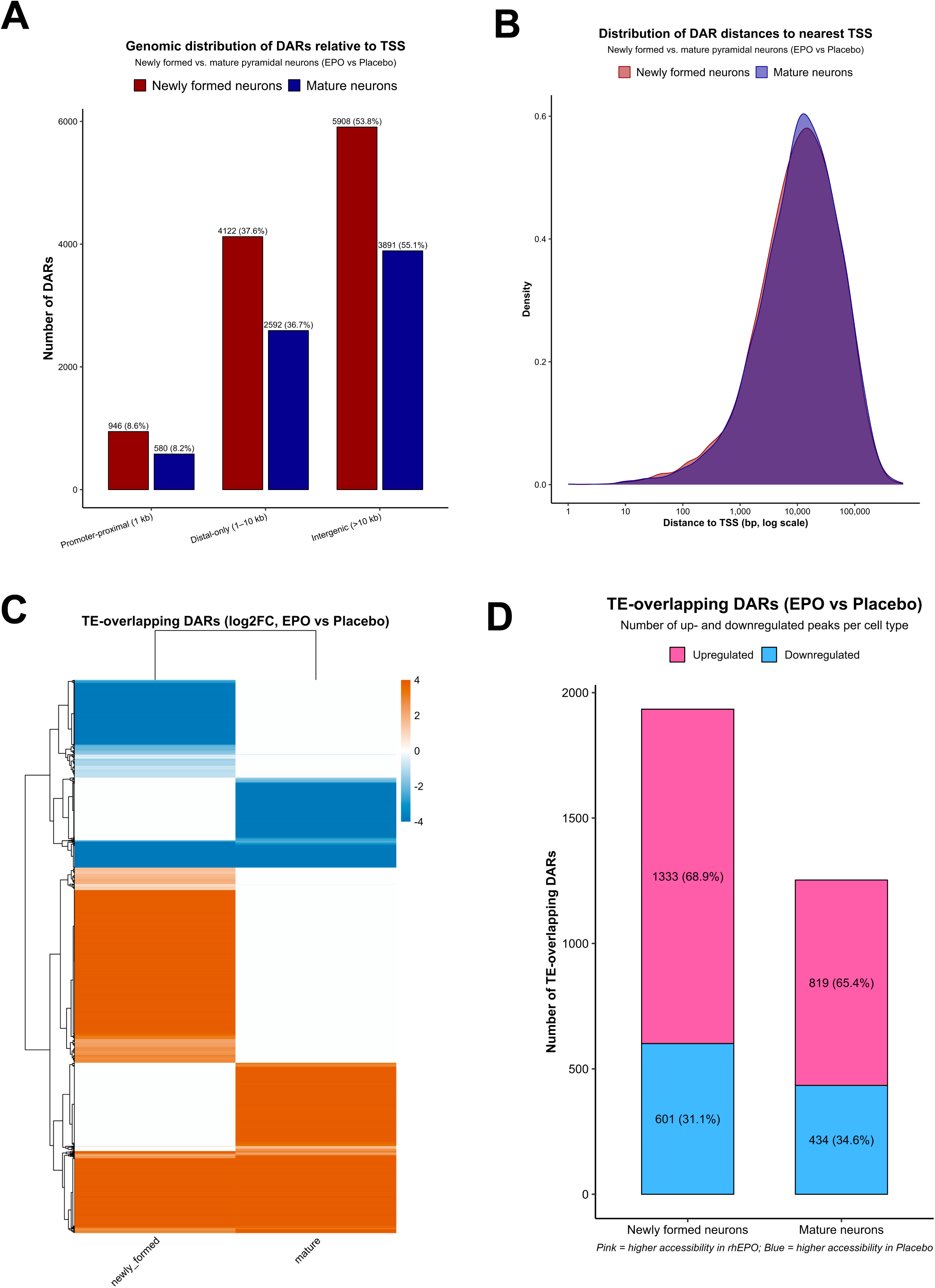
Genomic distribution and TE association of DARs in newly formed and mature pyramidal neurons. **(A)** Bar plot showing the number of DARs located in promoter-proximal (<=1 kb), distal (1-10 kb) and intergenic (>10 kb) regions for newly formed and mature pyramidal neurons (rhEPO versus placebo). (**B)** Density plot of distances from DAR summits to the nearest TSS on a log-scaled x-axis. **(C)** Heatmap of log2 fold-change values for TE-overlapping DARs in newly formed and mature neurons. **(D)** Bar plot summarising the number of TE-overlapping DARs that increase (pink) or decrease (blue) in accessibility in response to rhEPO within newly formed and mature neurons.

**Extended Data Fig. S12.**
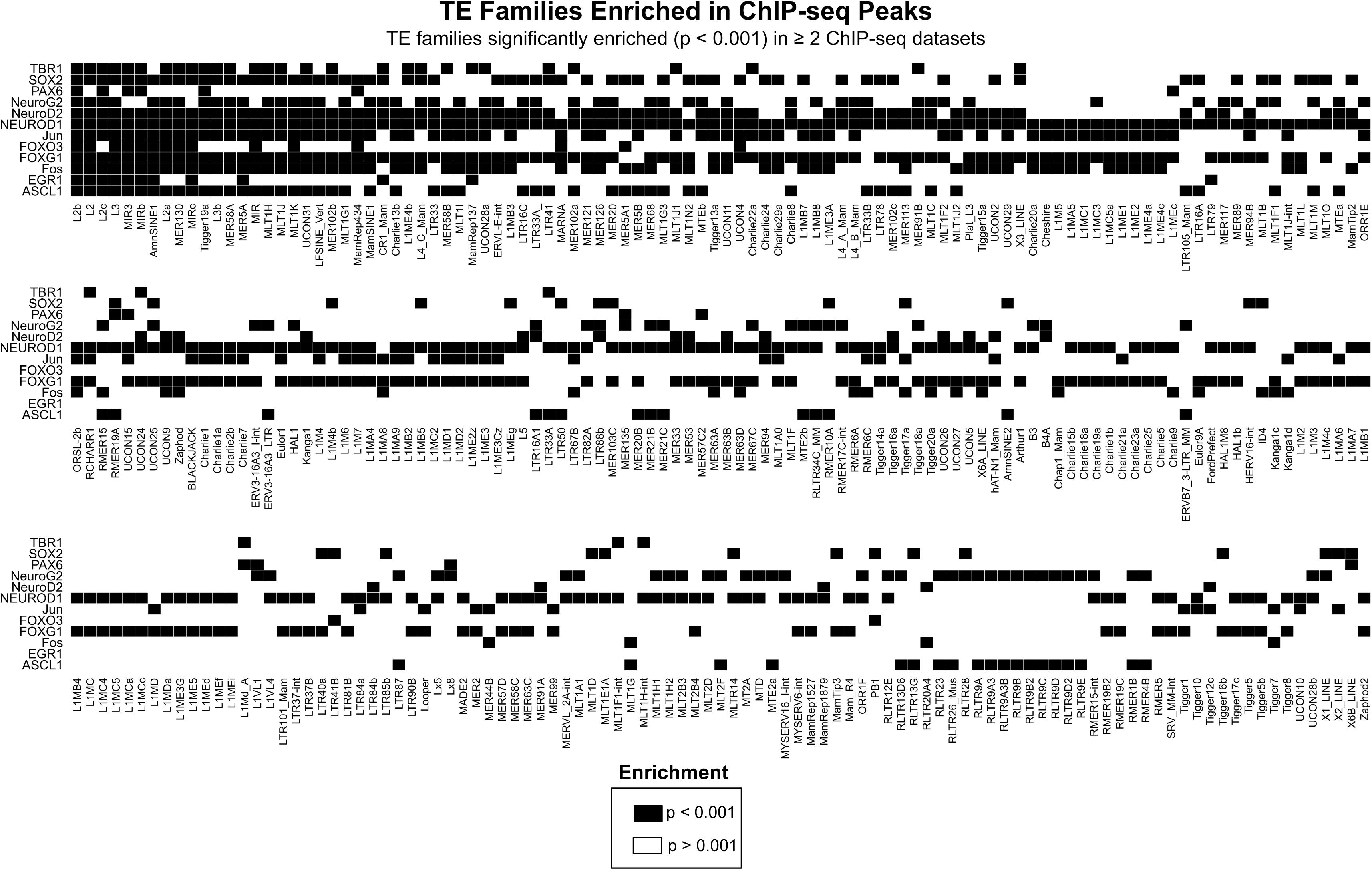
Transposable element families enriched within neurogenic transcription factor ChIP-seq peaks. Matrix showing TE families significantly enriched (p < 0.001) within ChIP-seq binding sites of neurogenic transcription factors. Rows correspond to transcription factors and columns to individual TE families; black squares indicate significant enrichment and white squares non-significant enrichment. Only TE families enriched in >= 2 independent ChIP-seq datasets are shown.

**Extended Data Fig. S13.**
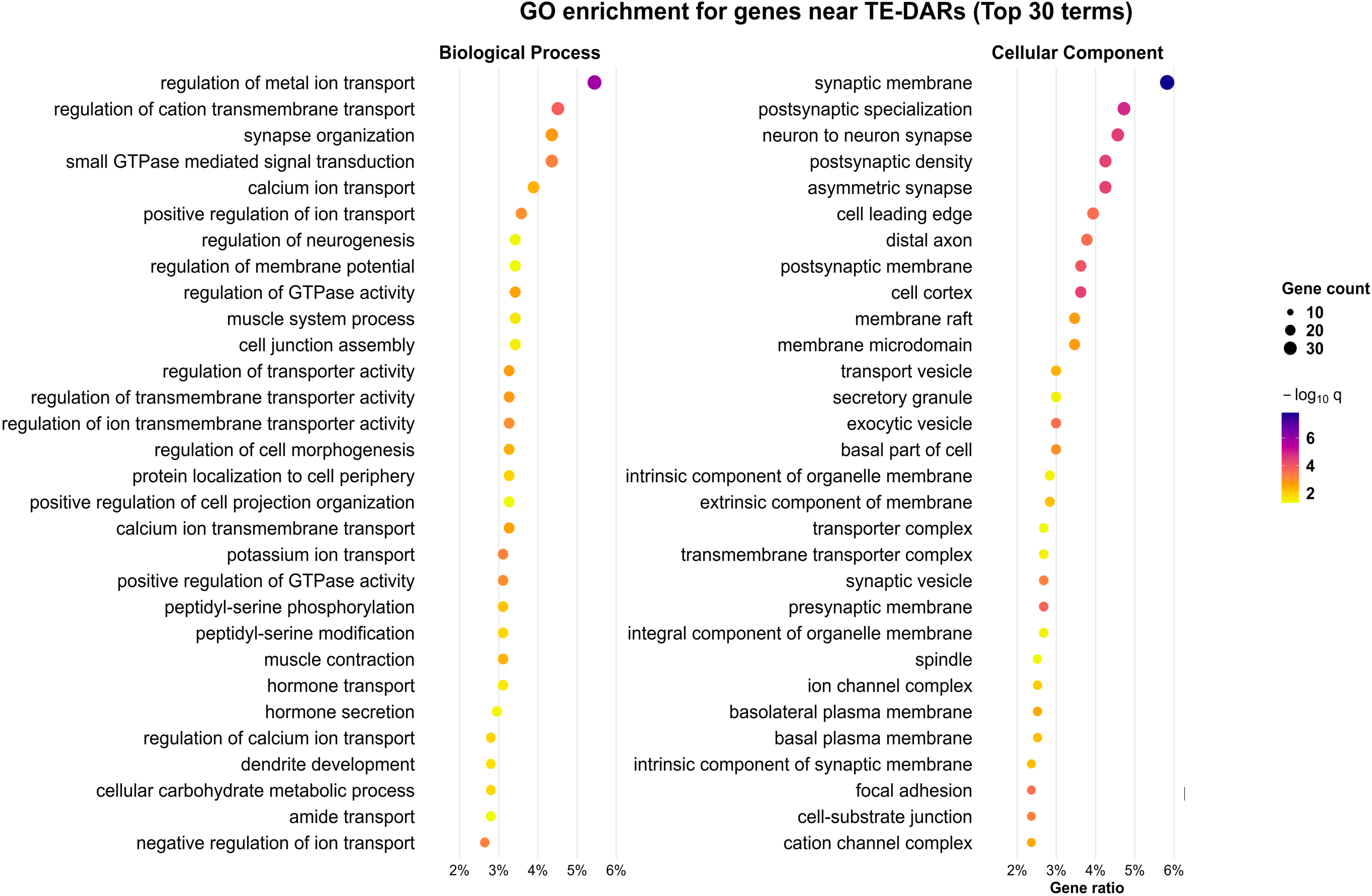
Gene ontology enrichment for genes located within +/-10 kb of TE-overlapping DARs. Dot plots showing the top 30 enriched Gene Ontology terms for genes whose transcription start sites lie within +/-10 kb of TE-overlapping DARs (rhEPO versus placebo), shown for Biological Process and Cellular Component. Dot size indicates the number of genes associated with each term and dot colour the statistical significance (-log10 q-value).

## SUPPLEMENTARY INFORMATION

⍰ Table S1: Summary of sequencing and alignment metrics for snATAC-seq samples, including per-sample nucleus counts.
⍰ Table S2: Transcription factor motif enrichment across Seurat-defined hippocampal clusters without treatment separation.
⍰ Table S3: Transcription factor motif enrichment across the 11 major hippocampal lineages without treatment separation.
⍰ Table S4: Summary of significant accessible chromatin peaks across the 11 hippocampal lineages and annotation of peaks not present in the Mouse Brain Atlas.
⍰ Table S5: Transposable element family enrichment within all accessible chromatin regions.
⍰ Table S6: Globally defined differentially accessible regions, motif enrichments, TE-associated regulatory elements and matched snRNA-seq comparisons for rhEPO and placebo conditions.
⍰ Table S7: Correlation between differentially accessible regions and differentially expressed genes after filtering for significant values.
⍰ Table S8: Significantly differentially accessible regions between rhEPO- and placebo-treated samples across the 11 hippocampal lineages.
⍰ Table S9: Significantly differentially accessible regions between rhEPO- and placebo-treated samples across the 11 hippocampal lineages, annotated for proximal genes and overlapping transposable elements.
⍰ Table S10: Gene regulatory network modules inferred for each of the 11 hippocampal lineages under rhEPO and placebo conditions.
⍰ Table S11: Enrichment of transposable element families within differentially accessible regions identified between rhEPO- and placebo-treated samples.
⍰ Table S12: Significant accessible chromatin peaks distinguishing newly formed from mature pyramidal neuron lineages.
⍰ Table S13: Differentially accessible regions between rhEPO- and placebo-treated samples within newly formed and mature pyramidal neurons, annotated for nearby genes and transposable elements.
⍰ Table S14: Enrichment of transposable element families within ChIP-seq peak sets for neurodevelopmental transcription factors.
⍰ Table S15: Transposable element-associated differentially accessible regions located within 10 kb of annotated transcription start sites.
⍰ Table S16: Gene Ontology analysis of genes associated with TE-overlapping differentially accessible regions bound by transcription factors.
⍰ Table S17: List of neurogenic transcription factors included in TF-binding and enrichment analyses.

## Notes

https://github.com/umutcakir/ATAC_EPO_vs_Placebo

https://doi.org/10.5281/zenodo.17635636

### Competing Interest Statement

The authors have declared no competing interest.

